# Enrichment of hard sweeps on the X chromosome compared to autosomes in six *Drosophila* species

**DOI:** 10.1101/2023.06.21.545888

**Authors:** Mariana Harris, Bernard Kim, Nandita Garud

## Abstract

The X chromosome, being hemizygous in males, is exposed one third of the time increasing the visibility of new mutations to natural selection, potentially leading to different evolutionary dynamics than autosomes. Recently, we found an enrichment of hard selective sweeps over soft selective sweeps on the X chromosome relative to the autosomes in a North American population of *Drosophila melanogaster*. To understand whether this enrichment is a universal feature of evolution on the X chromosome, we analyze diversity patterns across six commonly studied *Drosophila* species. We find an increased proportion of regions with steep reductions in diversity and elevated homozygosity on the X chromosome compared to autosomes. To assess if these signatures are consistent with positive selection, we simulate a wide variety of evolutionary scenarios spanning variations in demography, mutation rate, recombination rate, background selection, hard sweeps, and soft sweeps, and find that the diversity patterns observed on the X are most consistent with hard sweeps. Our findings highlight the importance of sex chromosomes in driving evolutionary processes and suggest that hard sweeps have played a significant role in shaping diversity patterns on the X chromosome across multiple *Drosophila* species.

## Introduction

The X chromosome has long been a subject of substantial interest in evolutionary biology due to its unique features that set it apart from the autosomes. Notably, the X harbors genes responsible for speciation, fertility, sexual dimorphism, and brain function (Rice 1984; Turelli and Orr 1995; Coyne 1998; Saifi and Chandra 1999; Skuse 2005; Payseur *et al*. 2018), highlighting its biological importance. Moreover, previous work suggests that the X chromosome may serve as a potential target of sexually antagonistic selection (Dean and Mank 2014; Patten 2019; Glaser-Schmitt *et al*. 2021), further emphasizing its significance in evolution. Consequently, studying adaptation on the

X chromosome, and how it differs from that of autosomes, can provide insights into the mechanisms driving genetic diversity, sexual selection, and speciation, thereby deepening our understanding of the broader processes that shape genetic variation across populations.

Adaptation on the X may differ from that of the autosomes due to two key differences. First, the X is expected to have a lower effective population size (*N_eX_*) compared to autosomes (*N_eA_*), leading to a decreased influx of new mutations. Second, due to male hemizygosity, new mutations on the X of males are immediately exposed to natural selection. This increased exposure to selection may lead to a higher probability of fixation of new recessive beneficial mutations (“Faster-X” effect (Charlesworth *et al*. 1987)) and a more efficient purging of deleterious variation on the X compared to autosomes, leading to lower levels of standing genetic variation on the X. Thus, as a consequence of these two factors, at the onset of positive selection, there will be a lower adaptive mutational supply on the X, resulting in more gradual rates of adaptation, or in other words, fewer haplotypes rising to high frequency bearing the adaptive allele (Orr and Betancourt 2001; Vicoso and Charlesworth 2006, 2009; Charlesworth *et al*. 2018).

Adaptation leaves behind distinct signatures in the genome. The classic signature, referred to as a hard selective sweep, occurs when a single adaptive mutation rises in frequency, resulting in deep dips in diversity in the vicinity of the adaptive locus (Maynard Smith and Haigh 1974; Kaplan *et al*. 1989). By contrast, a different signature known as a soft selective sweep, occurs when multiple adaptive mutations on distinct haplotypes sweep through the population simultaneously, not necessarily causing dips in diversity (Hermisson and Pennings 2005, 2017; Pennings and Hermisson 2006a; Messer and Petrov 2013). Recently, we found evidence of an enrichment of hard sweeps on the X chromosome compared to autosomes in a North American population of *D. melanogaster* (Harris and Garud 2023), suggesting that the X chromosome is subject to different evolutionary dynamics than the autosomes. Whether enrichment of hard sweeps on the X is a universal feature of molecular evolution has yet to be determined, as only a few species have been shown to have a higher prevalence of hard sweeps on the X compared to the autosomes (Nam *et al*. 2015; Harris and Garud 2023). Quantifying the prevalence of hard versus soft sweeps in natural populations has been of great interest and debate (Peter *et al*. 2012; Assaf *et al*. 2015; Schrider *et al*. 2015; Schrider and Kern 2016; Harris *et al*. 2018b; Feder *et al*. 2021; Garud *et al*. 2021). Thus, understanding the prevalence of hard and soft sweeps more broadly is crucial as it can shed light on common mechanisms that underlie adaptation in natural populations.

To understand if the X is generically enriched for hard sweeps in many species, we analyze population genomic data from six *Drosophila* species. By leveraging whole genome samples from multiple species and populations (Arbiza *et al*. 2014; Nam *et al*. 2015; McGrath 2022), we can identify trends that are the norm across species, as well as exceptions that are indicative of the unique biology of individual species. However, four of the populations we analyze in this study have small sample sizes (*n*=7-23 samples), making it difficult to conduct the same haplotype-based scan (Garud *et al*. 2015; Harris and Garud 2023) used in our previous work (Harris and Garud 2023). Additionally, each of these species has a unique demographic history for which we do not have accurate models, rendering our previous simulation-based approach for classifying putative sweeps as hard and soft challenging (Harris and Garud 2023). To overcome these challenges, we use a combination of single-nucleotide diversity and haplotype homozygosity statistics to analyze the patterns of diversity on the autosomes and the X chromosome across species, specifically looking for evidence of hard sweeps on the X that is inconsistent with other neutral and selective forces including background selection (BGS) and soft selective sweeps. Our empirical and simulation analyses show evidence that hard sweeps have played a significant role in shaping diversity patterns on the X chromosome in multiple *Drosophila* species suggesting that hard sweeps on the X are the norm rather than the exception.

## Methods

### Data

We analyzed data from six *Drosophila* species from seven populations (*D. melanogaster* from Zambia (ZI), *D. melanogaster* from Raleigh (RA)*, D. simulans, D. sechellia, D. mauritiana, D. santomea* and *D. teissieri*), which we downloaded and processed as follows: For both *D. melanogaster* populations, data are publicly available as part of the Drosophila Genome Nexus data set (Lack *et al*. 2015) and can be downloaded from www.johnpool.net. We downloaded 205 DGRP genomes from Raleigh (RA), North Carolina and 197 DPGP3 genomes from Zambia (ZI). For our analysis we used 100 genomes from each population, previously processed (Harris and Garud 2023) to remove individuals having high IBD with one another, as well as residual heterozygosity, and high levels of missing data.

For *D. simulans* we downloaded data from 170 inbred lines from a North American population (Signor *et al*. 2018) available at https://zenodo.org/record/154261#.YzMzty2z3jC. For the remaining four species, we obtained genomes from NCBI’s RefSeq and short read data from NCBI’s Short Read Archive (Garrigan *et al*. 2014; Turissini and Matute 2017; Meany *et al*. 2019; Serrato-Capuchina *et al*. 2021) (**Table S1**).

For *D. mauritiana, D.sechellia, D. teissieri* and *D. santomea* variant calling was performed with the NVIDIA Clara Parabricks pipeline v4.0.0 (https://docs.nvidia.com/clara/parabricks/4.0.0/index.html), a re-implementation of variant calling tools including BWA-mem (Li 2013) and GATK4 (Van der Auwera and O’Connor 2020) optimized for running on NVIDIA graphics processing units. Following the GATK best practices for germline variant calling (https://gatk.broadinstitute.org/hc/en-us/sections/360007226651), we mapped each sample’s reads to the appropriate reference genome, sorted and removed PCR duplicates, generated single-sample GVCFs, then performed joint genotyping with HaplotypeCaller. From this initial set of variant calls, we removed sites with quality scores QUAL<30.0 to obtain a bootstrap set of high-confidence calls for another round of variant calling (https://gatk.broadinstitute.org/hc/en-us/articles/360035890531-Base-Quality-Score-Recalibration-BQSR). We performed base quality score recalibration on the mapped reads using the bootstrap set then once again generated single sample GVCFs that were used for joint genotyping with HaplotypeCaller to ultimately generate a VCF file for each species.

To remove any low quality variant sites, we applied GATK recommended hard filters (https://gatk.broadinstitute.org/hc/en-us/articles/360035890471-Hard-filtering-germline-short-variants), summarized as follows: FS > 60, QD < 2.0, MQ < 4, MQRankSum < -12.5, QUAL < 30.0, SOR > 3.0, and ReadPosRankSum < -8.0. Additionally, we excluded sites that were not uniquely mappable (**Fig. S1**). We also filtered out sites lying within repetitive elements as predicted by RepeatMasker (Smit *et al*. 2015). Only bi-allelic SNPs were considered in our analysis.

Next, we excluded invariant sites of poor quality. To do so we filtered entire regions, which included both variant and invariant sites, with poor depth and quality statistics. We used banded gVCF files, where sites of similar quality get concatenated into a band. We excluded intervals that failed to meet the depth and quality criteria: (’MIN(FMT/DP)>10 & MIN(FMT/GQ)>30’). Next, we combined the results from the GATK hard filters applied to variant sites with the resulting regions – comprising both variant and invariant sites – that passed the filter applied to the banded gVFC files. This gave us the total number of callable sites in the genome. When computing statistics using a sliding window approach (see section below), we excluded windows that overlapped 50% more with low-quality intervals.

High-density repeat regions are often associated with centromeric regions, which consist of highly homogeneous tandem repeats and are known to experience low rates of recombination (Mather 1939; Levine 1955; Vincenten *et al*. 2015). Consequently, these regions tend to exhibit reduced diversity and increased homozygosity, which can lead to false positive signals of selection. To mitigate potential confounding effects of repeats on selection inferences, we calculated the proportion of base pairs identified as repeats within 50Kb windows. In addition to removing individual sites masked as repeats, we removed entire windows in which 20% or more of the sites were marked as repeats by RepeatMasker.

### Diversity statistics and haplotype homozygosity measured from data

We annotated which SNPs lie in exon, intron or intergenic regions using RefSeq (O’Leary *et al*. 2016) annotations. The corresponding assembly accession numbers for each species are: GCF_004382145.1 (*D. mauritiana)*, GCF_004382195.2 (*D. sechellia*), GCF_016746235.2 (*D. teissieri*), GCF_016746245.2 (*D. santomea*) and GCF_000001215.2 (*D. melanogaster*). For *D. simulans* we used the reference genome and corresponding gff file provided by Rebekah Rogers and Peter Andolfatto (Rogers *et al*. 2014). We used BEDtools v.2.3.0 (Quinlan and Hall 2010) to separate the data into exon, introns or intergenic regions. If any position could be included in multiple categories, we annotated the position prioritizing exons followed by introns and finally intergenic regions. Additionally, we excluded regions less than 40bp long.

To understand the influence of selection across genomic regions on the X chromosome versus autosomes, we calculated nucleotide diversity in autosomes (π_A_) and the X chromosome (π_X_) in each of the exons, introns, and intergenic regions. We next computed the ratio of X to autosomal diversity (π_X_/π_A_) by running 1,000 bootstrap replicates in which we sampled exons, introns, and intergenic regions at random, computing the mean π_X_, mean π_A_, and mean(π_X_)/mean(π_A_) in each sample. We approximated π by π ≈ 2*pq*, with *p* the frequency of the major allele at a given site and *q*=1-*p*.

To identify regions of the genome that may have exceptionally low diversity due to selection, we also computed π/bp in non-overlapping windows of 10, 20 and 50 kb (**Fig. S2**) across the autosomes and the X chromosome of all species. We then defined a low diversity threshold as X% of the π/bp average of each chromosome with X=20 or 30%. We labeled windows with π/bp below this threshold as low diversity windows, further investigated as putative sweep regions. For the main analysis of this work, 20Kb windows and a threshold of 20% of the average π/bp was used. To test for differences in quality in windows below and above the defined threshold, we computed the proportion of missing data and quality score per site and found similar distributions across these categories (**Fig. S3**).

Additionally, we computed haplotype or multi-locus genotype homozygosity in windows of 20 SNPs long on the X chromosome, with the expectation that haplotype homozygosity should be elevated in selective sweeps but not background selection (Wall and Pritchard 2003; Enard *et al*. 2014; Garud *et al*. 2015; Schrider 2020). In species for which we had phased data (*D. melanogaster* and *D. simulans*) we computed the expected haplotype homozygosity (Garud *et al*. 2015) defined as 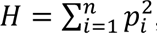, where *p*_*i*_ is the frequency of the *i^th^* most common haplotype in a sample with *n* distinct haplotypes. For the remaining species we computed the expected multi- locus genotype homozygosity (Harris *et al*. 2018a) defined as 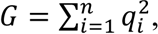, where *q*_*i*_ is the frequency of the *i^th^* most common multi-locus genotype in a sample with *n* distinct haplotypes. In contrast to a phased haplotype, where the allelic state for each site is known, a multi-locus genotype is a string that represents the diploid state of the individual where each site is labeled as either homozygous for the reference allele, homozygous for the alternate allele, or heterozygous. A high recombination rate is expected to break haplotypes and thus decrease both H and G, whereas with a low recombination rate, haplotypes may drift to high frequency and thus increase H, and by extension G.

In addition to measuring H and G in 20 SNP windows, we tested windows of length 50 and 10 SNPs long. However, due to the small sample size of most species, the probability of observing the same multi-locus genotype twice becomes small with longer window sizes, making it difficult to capture any signal in the 50 SNP window case. With 10 SNP windows we also found an elevation of homozygosity in low diversity windows. However, we opted for 20 SNP windows as smaller window sizes may increase homozygosity due to genetic drift.

### Simulation analysis

To understand the evolutionary processes responsible for the patterns of π/bp and haplotype homozygosity observed in the data, we simulated a variety of evolutionary models for the X chromosome and autosomes using SLiM 3.7 (Haller and Messer 2019). The scenarios simulated included neutrality, sex bias, low recombination rate, mutation rate bias, bottlenecks, background selection, hard sweeps and soft sweeps as described below.

For all models with a fixed population size (excluding bottleneck models) we simulate a constant *Ne*=10^6^ population. An *N_e_* of this order of magnitude is reasonable for most of the species analyzed given mean nucleotide diversity levels, with exception of *D. sechellia*, which has been shown to have a lower *Ne*∼10^5^ (Legrand *et al*. 2009). SLiM is a forward in time simulator, which makes simulating populations with *N_e_* > 5x10^5^ unfeasible due to memory requirements. As in our previous work (Harris and Garud 2023) we rescaled our simulations using a constant factor of Q=50.

In all our simulations we modeled a 20kb region with a recombination rate of *r* =5x10^-7^ cM/bp (unless otherwise specified) and a neutral mutation rate of µ=1x10^-9^, both rescaled by Q=50. For the simulations that include selection we assumed that mutations on the X of males experience the same fitness effect as that of a homozygous female (i.e. dosage compensation). We ran a total of 100 simulations for each model and ran a *10N_e_* burn-in for every simulation.

### Neutral models

We first simulated a completely neutral model with a female to male ratio of one and equal mutation rates between the sexes. Next, we introduced scenarios that can differentially impact X-linked and autosomal diversity. We simulated female and male sex bias varying the sex ratio as 2:1, 5:1 and 7:1 for each scenario, as well as a model with a lower X-linked mutation rate reducing the X-linked mutation rate such that the ratio µ_X_/µ_A_ was equal to 0.1,0.5, 0.75 or 0.9. We also tested the effect of regions of low recombination by running simulations with recombination reduced to 1,10, 20 and 50% of the original *r*.

The expected lower population size of the X can result in stronger drift that can be exacerbated in a bottleneck. For this reason, we considered two bottleneck models that were fit to to π/bp and S/bp in short introns from DGRP dataset (Garud *et al*. 2015, 2021): (i) a severe and short bottleneck with a bottleneck *N_e_* of 0.002*N_e,ancestral_* for 0.0002*\*2N_e,ancestral_* generations, and (ii) a shallow and long bottleneck with a bottleneck *N_e_* of 0.4*N_e,ancestral_* for 0.056*\*2N_e,ancestral_* generations.

### Background selection

Background selection can result in stronger dips in diversity on the X compared to autosomes (Charlesworth *et al*. 1993, 1995; Stephan 2010). To consider this scenario, we simulated a constant *Ne*=10^6^ model with deleterious variation. The selection coefficients for deleterious mutations were gamma distributed with mean and shape parameter -0.000133 and 0.35, respectively (Huber *et al*. 2017). We varied the percentage of deleterious mutations simulating the case in which 10, 50 or 80% of incoming mutations were deleterious. Additionally, we simulated BGS and varied the X-linked mutation rate in both bottleneck models described previously to test whether the effect of both processes can produce the patterns in the data. We did this for *r*=2.5e-7 cM/bp and *r*=5e-7 cM/bp. Furthermore, we simulated 1Mb chromosomes varying the recombination rate (r=0, 1e-8, 5e-7 cM/bp) to assess the effect of a higher mutation load and recombination rate in the context of BGS.

### Selective sweeps

To understand the effects of positive selection on patterns of diversity on the X chromosome and autosomes we simulated hard and soft sweeps. To model hard sweeps, we introduced a single adaptive mutation on the center of the haplotype (θ_a_=0.01) and restarted the simulation if the adaptive mutation was lost. We conditioned on fixation of the sweep and sampled 100 haplotypes.

To simulate soft sweeps, we introduced adaptive mutations recurrently at the center of the haplotype at a rate defined by θ_a_=0.1, 1 and 10, where θ_a_=4*Neµ_a_* with *µ_a_* the adaptive mutation rate. We conditioned our simulations on the fixation of the sweep, after which we took a sample of n=100 haplotypes. We verified that our simulation represented a soft sweep by only including samples with two or more mutational origins in our analysis. For both hard and soft sweeps, we varied the strength of selection such that *Nes_b_*=20, 200 or 2000.

### Identification of shared genes under selection in multiple species

To investigate whether similar functions are selected for on the X in multiple species, we obtained the genes intersecting windows below and above the low diversity threshold using BEDtools v.2.3.0 and the annotation files from the reference genome for each species in our study. Next, we used the gene IDs and matched them to their corresponding ortholog IDs in orthoDB v11 (Kuznetsov *et al*. 2023). We then looked for overlapping orthologs across species and obtained the proportion of orthologs that overlap over two, three or four species in low versus high diversity windows.

To understand whether there were more shared regions under selection in low versus high diversity regions across species, we obtained 100 random samples of *n* orthologs from high diversity regions with *n* the number of orthologs in low diversity regions per species. From this we computed the mean proportion of shared orthologs under selection as well as the 95% confidence interval in high diversity regions.

### Code availability

Code used for simulations as well as to process and analyze the data is available at https://github.com/garudlab/DrosCrossSpecies_XchrHardSweeps

## Results

We analyzed population genomic data of six different *Drosophila* species from seven populations, including: *D. melanogaster* (n=100 Zambia (ZI), n=100 Raleigh (RA))*, D. simulans* (n=170)*, D. sechellia* (n=23)*, D. mauritiana* (n=15)*, D. santomea* (n=7) and *D. teissieri* (n=11) (**Table S2**). To understand not only how selection varies across species, but also across populations within a species, we include a derived *D. melanogaster* population from Raleigh as well as a population from Zambia that is presumed to be within the ancestral range of the species (Pool *et al*. 2012). First, we compare the patterns of nucleotide diversity (π) between the autosomes and the X chromosome in the data. Next, we analyze haplotype homozygosity in low diversity regions on the X compared to the rest of the chromosome. Finally, we simulate a variety of models, both without and with selection, to test whether any of these can generate the patterns observed in the data.

### Patterns of X versus autosomal diversity across the genomes of six *Drosophila* species

When a population is not subject to any selective forces, has the same mutation rate across sexes, a male-to-female ratio of one, and a constant effective population size (*N_e_*), the diversity on the X chromosome (π_X_) is expected to be 3/4 of the autosomal diversity (π_A_) (Vicoso and Charlesworth 2006). However, such a simple model rarely captures the complex dynamics of natural populations, rather, it is likely that different evolutionary forces affect the X and autosome differently, leading to deviations from the π_X_/π_A_=0.75 expectation. Across the seven populations studied, we observed a significantly lower diversity on the X compared to autosomes with five out of seven populations showing π_X_/π_A_ below 0.75 (**Fig. 1**). The other two populations (*D. melanogaster* (ZI) and *D. teissieri*) also deviated from the π_X_/π_A_=0.75 expectation but in the opposite direction, with π_X_/π_A_>0.75. This trend has been previously reported for African populations of *D. melanogaster* and has been attributed to other evolutionary forces such as female sex bias (Kauer *et al*. 2002; Singh *et al*. 2007; Pool *et al*. 2012).

**Figure 1.**
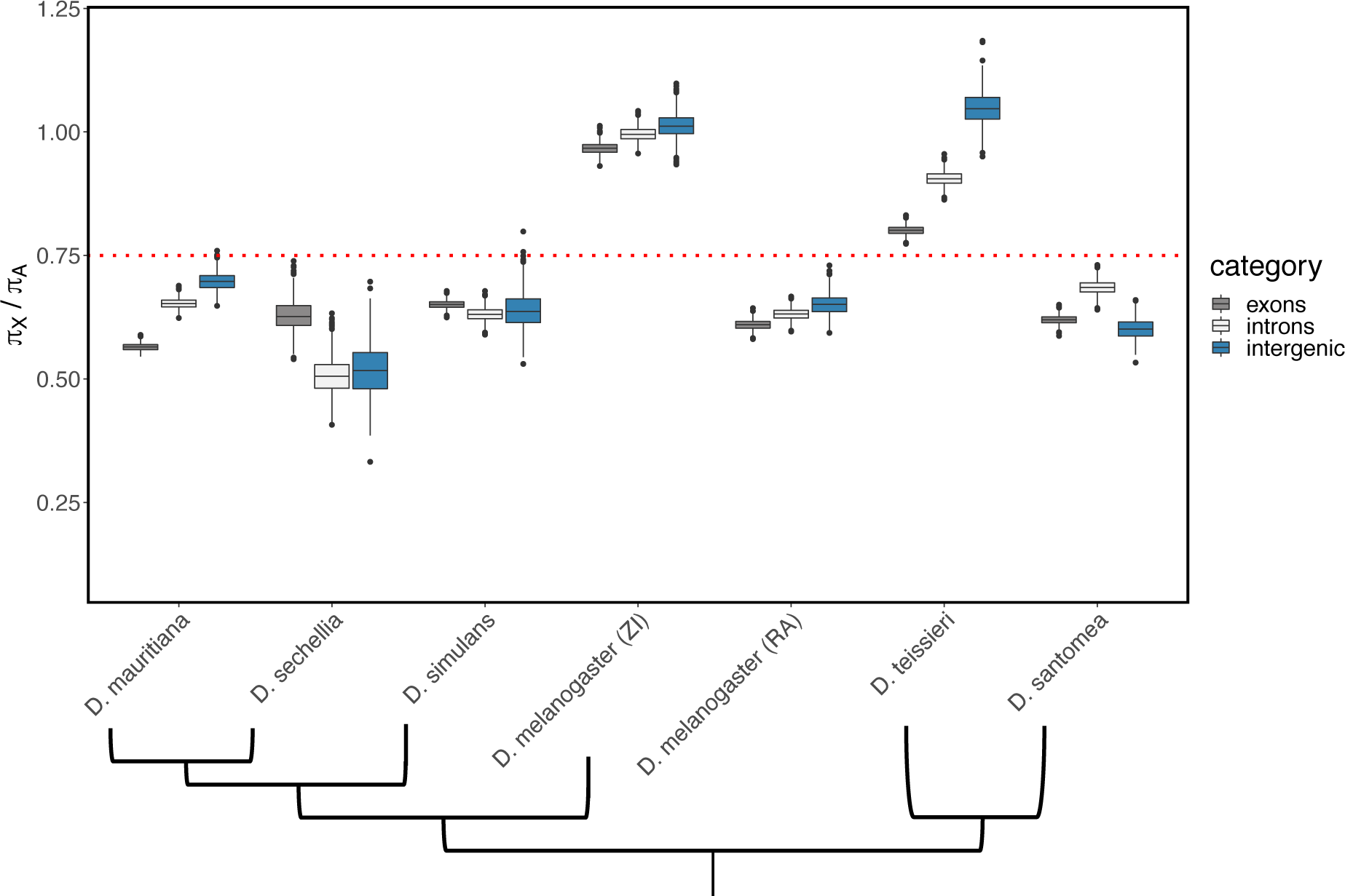
Nucleotide diversity (π/bp) in exons, introns and intergenic regions on the autosomes and the X chromosome of six *Drosophila* species. π_X_/π_A_ is lower on exons than on introns and intergenic regions in most species. The red dotted line corresponds to the 0.75 expectation under a model with an unbiased sex ration and equal mutation rates on the X and autosomes. For each category we performed 1,000 bootstrap replicates (**Methods**). The tree shown below the X-axis represents the phylogenetic relationship between species as described in FlyBase (Gramates *et al*. 2022).

Functionally important regions of the genome, such as exons, are expected to be subject to selection, positive or negative (McVicker *et al*. 2009; Arbiza *et al*. 2014; Nam *et al*. 2015). Hence, to investigate the influence of selection across species, we divided the data into three regions: exons, introns, and intergenic regions and computed the ratio of X to autosomal diversity (π_X_/π_A_) for each region. We found a stronger reduction in diversity in exons compared to introns and intergenic regions for most species (**Fig.1**), pointing to the functional importance of exons (Nam *et al*. 2015). Moreover, we found that π_X_/π_A_ was consistently below 0.75 for all regions for most species (**Fig. 1**), suggesting more directional selection on the X compared to autosomes. However, we noted an exception in *D. sechellia,* where exons exhibited higher levels of diversity than introns and intergenic regions. This behavior could be explained by the high levels of introgression from *D. simulans* to *D. sechellia* (Garrigan *et al*. 2012; Matute and Ayroles 2014; Schrider *et al*. 2018). Introgression tends to be higher in non-functional regions of the genome and is generally higher on the autosomes than on the X due to the involvement of sex chromosomes in hybrid incompatibilities (Muirhead and Presgraves 2016; Turissini and Matute 2017; Fraïsse and Sachdeva 2021). This could lead to a greater difference between X and autosomal diversity in intergenic regions compared to exons, potentially explaining the patterns observed in *D. sechellia*’s data.

To identify potential regions under selection, we calculated π/bp in 20Kb windows across each chromosome for every species. Next, we labeled the analysis windows as low diversity windows if they fell below a diversity threshold set as 20% of the chromosomal π/bp average. Importantly, we computed this threshold for each chromosome separately to normalize for variance in coalescence time across chromosomes. We found that, for all species, the X chromosome showed a higher proportion of windows below the low diversity threshold compared to the autosomes (**Fig. 2** and **Fig. S4**). Remarkably, these low diversity windows often exhibited π/bp values significantly lower than the defined threshold with values as low 0.5% of the chromosomal average (**Table S3; Fig. S5**). Furthermore, we observed multiple instances of consecutive low diversity windows extending up to 200Kb, shown as the yellow shaded regions of **Fig. 2A**. To validate our findings, we repeated this analysis using 10Kb and 50Kb analysis windows as well as two different low diversity thresholds. In all cases, we observed an increased proportion of low diversity windows on the X compared to autosomes (**Fig. S2**).

**Figure 2.**
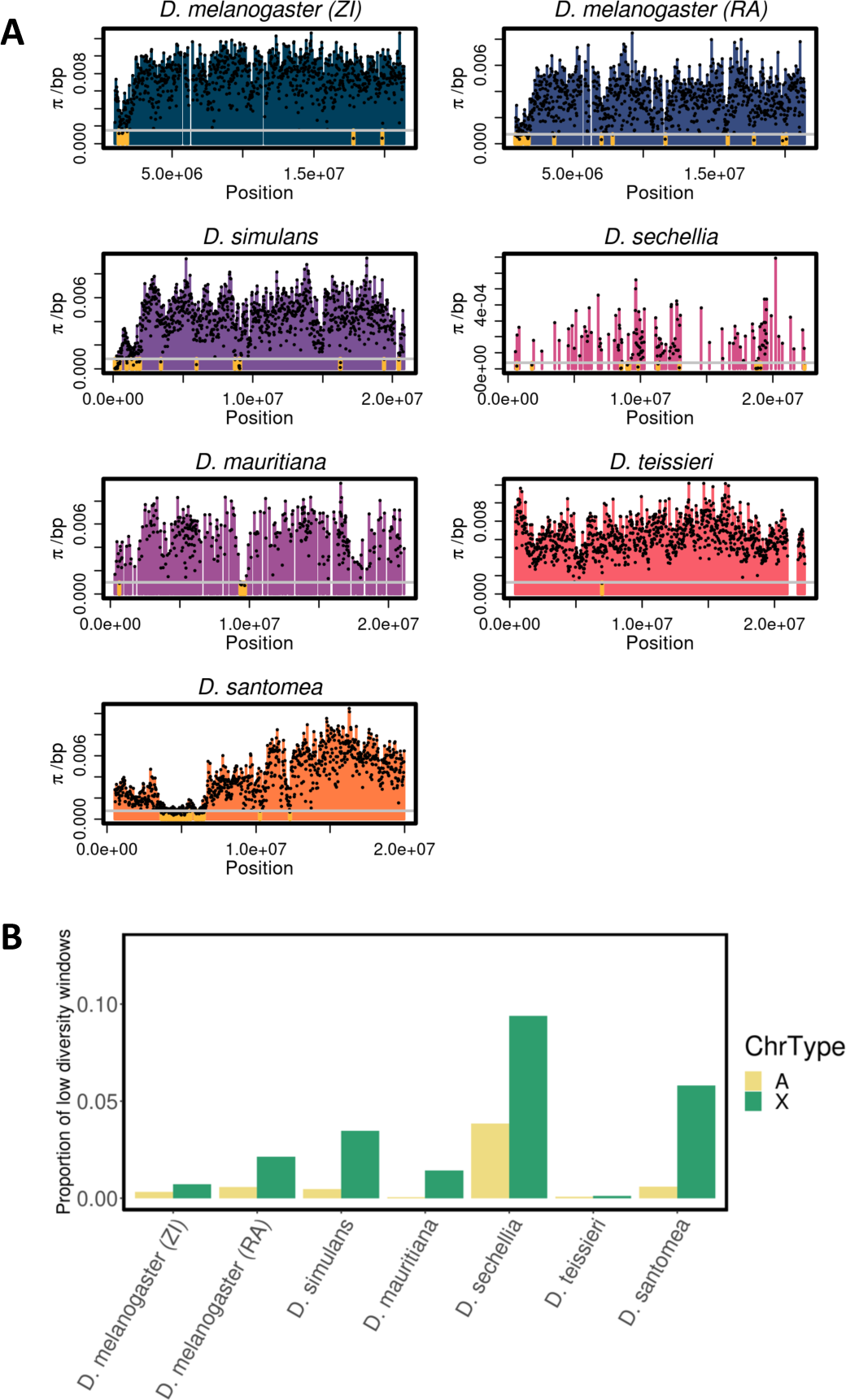
Multiple *Drosophila* species have a higher proportion of windows on the X with steep depressions of diversity compared to autosomes. (A) π/bp in 20kb windows on the X chromosome of six *Drosophila* species and two *D. melanogaster* populations from Raleigh (RA) and Zambia (ZI). Windows where diversity is below 20% of the chromosomal average (gray horizontal line) are highlighted in gold. In **Fig. S4**, the same figure for all autosomes is plotted for each species. **(B)** Proportion of windows below the 20% of the chromosomal average on the autosomes (yellow) and the X chromosome (green).

Elevated homozygosity can arise from different processes such as positive selection or population bottlenecks (Wall and Pritchard 2003). Therefore, to gain deeper insights into the potential mechanisms responsible for the dips in diversity on the X, we computed haplotype homozygosity (H) across windows below and above the low diversity threshold (**Methods**; **Fig. 3**) in species with phased genomes (*D. melanogaster, D. simulans* and *D. mauritiana*). For species with unphased genomes, we computed multilocus genotype identity (G) ((Harris *et al*. 2018a); **Methods**). We found that in all populations, windows below the low diversity threshold showed a significantly elevated homozygosity compared to the rest of the chromosome (one-sided Wilcoxon rank sum test p-val< 0.05; **Fig. 3**). Below we examine the role of selection in generating these differential patterns between the X and the autosomes.

**Figure 3.**
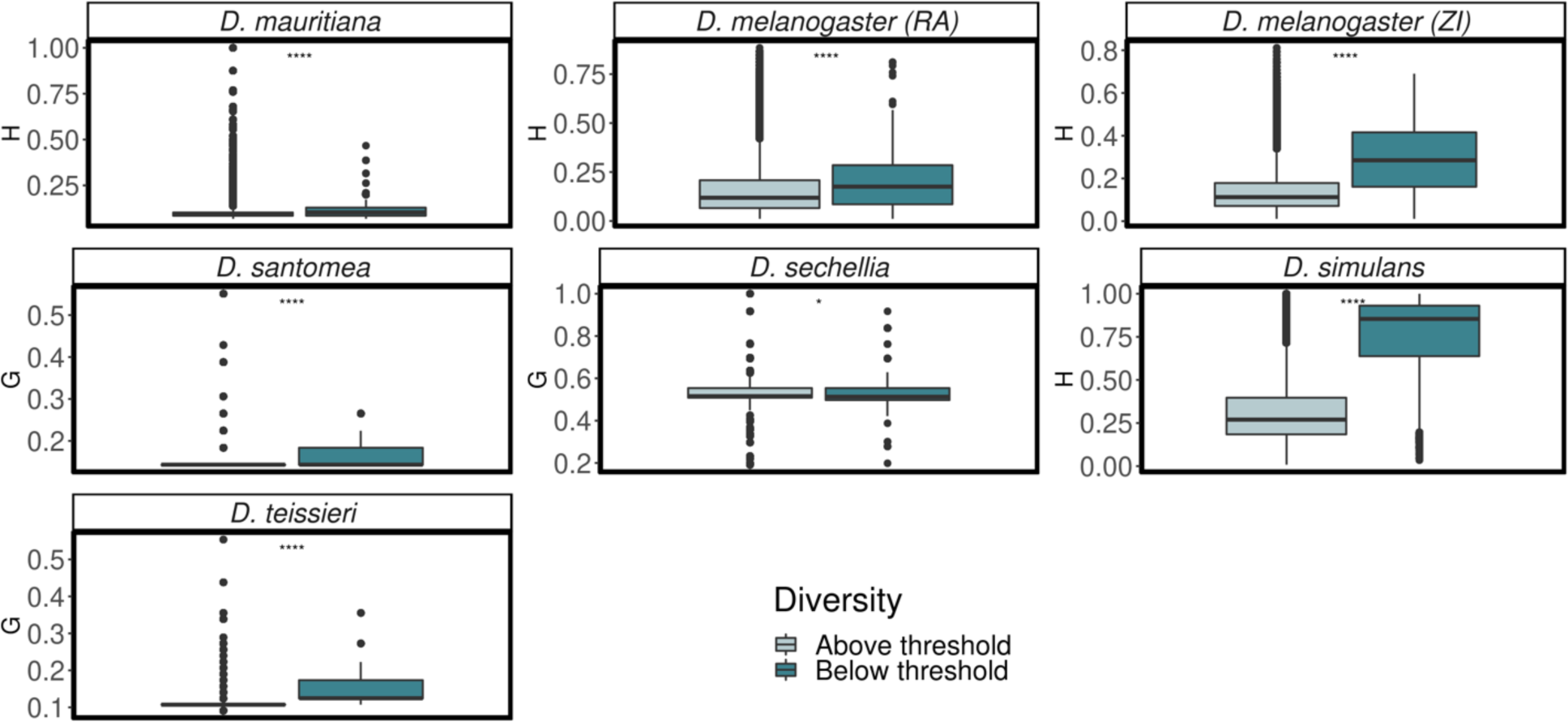
Homozygosity in windows below and above 20% of the average π/bp on the X chromosome. Windows with π/bp below 20% of the chromosomal mean show elevated haplotype homozygosity (H) or elevated multilocus genotype homozygosity (G) compared to windows with π/bp above this threshold (one-sided Wilcoxon rank sum test p-val< 0.05).

### Neutral models cannot recapitulate diversity patterns observed in the data

The differing diversity patterns on the X versus autosomes could be the result of distinct population genetic forces including: (1) sex biased demography, (2) differences in mutation rates between males and females, (3) lower recombination rates, (4) stronger effects of drift on the X given its smaller *N_e_* and/or (5) more efficient natural selection on the X due to male hemizygosity (Betancourt *et al*. 2004; Vicoso and Charlesworth 2009; Arbiza *et al*. 2014; Nam *et al*. 2015).

To gain a better understanding of how different evolutionary processes influence the relative X to autosomal diversity, we quantified π_X_, π_A_ and haplotype homozygosity across a wide variety of scenarios simulated with the population genetics simulator SLiM (Haller and Messer 2019). We started by simulating neutral models of sex bias, low recombination rates, mutation bias and population bottlenecks.

### Sex bias

Our simulations show that female bias can increase π_X_/π_A_ above the 0.75 expectation (**Fig. 4A**), recapitulating the patterns observed in the Zambian population of *D. melanogaster* and consistent with evidence of female bias in African *D. melanogaster* populations previously reported in the literature (Kauer *et al*. 2002; Dieringer *et al*. 2005; Thornton and Andolfatto 2006; Singh *et al*. 2007; Pool *et al*. 2012) **(Fig. 1** and **Fig. 4A)**. *D. teissieri* also has π_X_/π_A_>0.75, which could also be consistent with female bias. However, none of our female bias models produced elevated haplotype homozygosity compared to neutrality (**Fig. 4B**).

**Figure 4.**
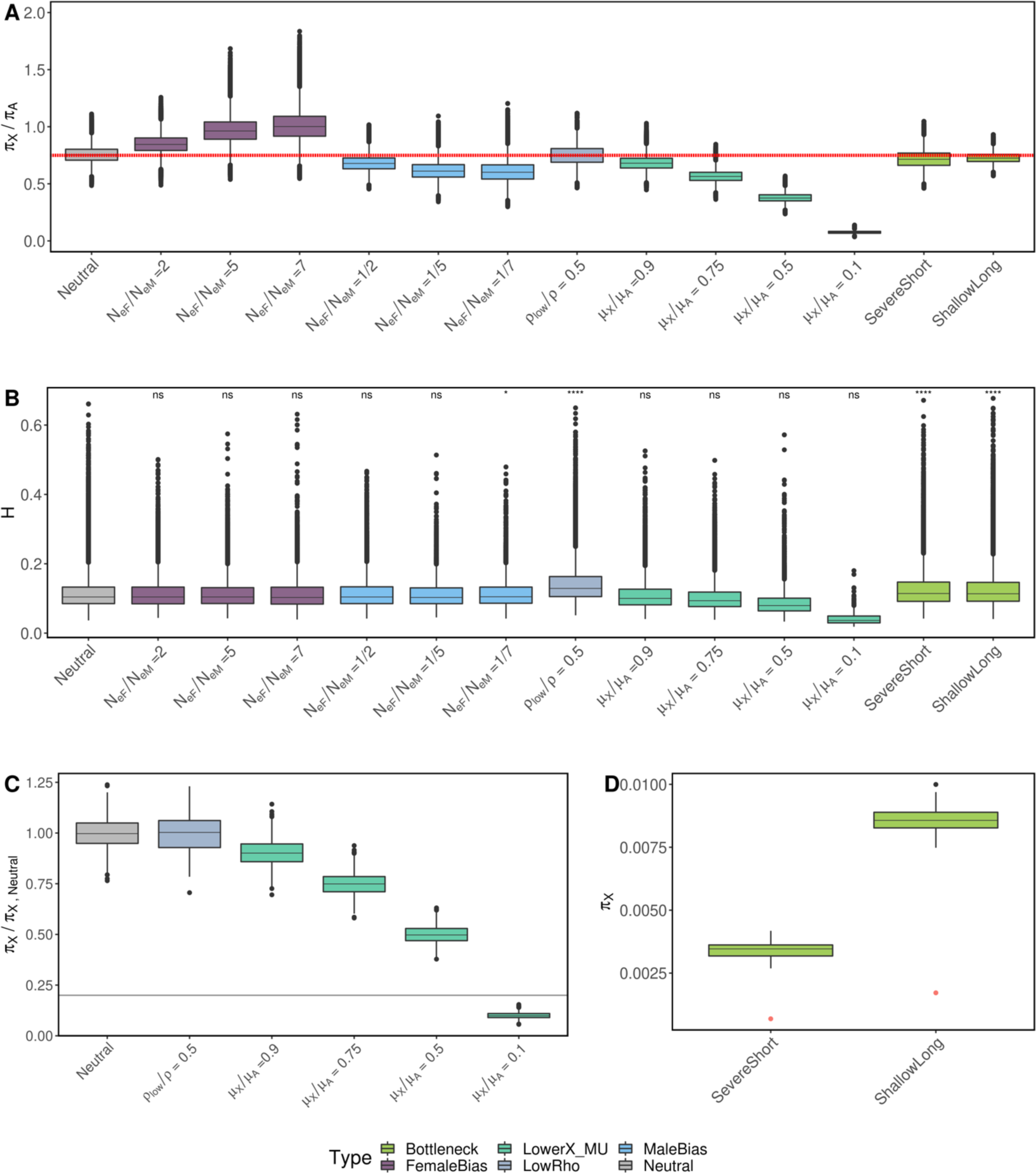
Nucleotide diversity (π) and haplotype homozygosity across 12 models with no selection. The models considered include three female-biased models (*N_eF_/N_eM_=*2, *N_eF_/N_eM_=*5, and *N_eF_/N_eM_=*7), three male-biased models (*N_eM_/N_eF_=*2, *N_eM_/_NeF_ =5*, and *N_eM_/_NeF_ =* 7), one low recombination model (*r_low_/r = 0.5*), four lower X-linked mutation rate models (µ_X_*/*µ_A_ =0.9, µ_X_*/*µ_A_=0.75, µ_X_*/*µ_A_=0.5, and µ_X_*/*µ_A_=0.1, where µ_X_ is the X-linked mutation rate and µ_A_ is the autosomal mutation rate), and two bottleneck models (a severe short bottleneck and a shallow and long bottleneck). **(A)** π_X_/π_A_ across all models, where the red dashed line corresponds to the expected π_X_/π_A_ = 0.75 value in the baseline neutral model (Same sex ratio, equal X and autosomal mutation rates and a constant *N_e_*=10^6^). **(B)** Haplotype homozygosity across all models. The asterisks represent a significant elevation in haplotype homozygosity relative to complete neutrality using a one-sided Wilcoxon rank-sum test. **(C)** π_X_/ π_X,Neutral_ across low X mutation rate models, with π_X,Neutral_ equal to the average π in the baseline neutral model. The horizontal gray line is the low diversity threshold, defined as 20% of the average π/bp of the neutral case. **(D)** π_X_ in two bottleneck models. The red data point below the boxplots corresponds to 20% of the average π from each bottleneck model.

Additionally, we found that male sex bias can decrease π_X_ such that π_X/_ π_A_ < 0.75, which could potentially explain low π_X_ values observed in some species (*D. melanogaster (RA), D. simulans, D. mauritiana* and *D. santomea;* **Fig. 1**). However, only the most extreme case of male bias (1:7 female-male bias) can marginally increase haplotype homozygosity compared to neutrality (**Fig. 4B**). To the best of our knowledge, such an extreme male-sex bias in the species studied has not been reported in the literature and therefore seems unlikely.

### Recombination rate variation

Another possibility is that regions with a low recombination rate produce dips in diversity and high haplotype homozygosity as observed in the data. The footprint of a hard selective sweep is inversely proportional to the recombination rate (∼ *s/r*), due to high recombination rates breaking linkage and impeding the formation of long haplotypes at high frequency (Maynard Smith and Haigh 1974; John H. Gillespie 2004). Consequently, it is reasonable to anticipate lower recombination rates in our putative hard sweep regions when compared to the overall genomic landscape. For *D. melanogaster*, where there is a detailed recombination rate available, we find that, on average, recombination rates are 37% and 48% lower in the low diversity regions of the RA and ZI populations, respectively (**Fig. S6**; (Comeron *et al*. 2012). To understand if a lower recombination rate can explain all the signatures in the data, we simulate windows with reduced recombination rates (*r_low_*), such that *r_low_ /r= 0.5, 0.2,0.1,0.01*. We found that while a low recombination rate can elevate haplotype homozygosity it cannot generate dips of diversity below the low diversity threshold (**Fig. 4, Fig. S7**).

### Mutation rates

Lower X-linked mutation rates could result in stronger dips in diversity on the X compared to autosomes. A lower X-linked mutation rate is expected when there is a higher number of germline cell divisions in males than females (Drost and Lee 1995; Kirkpatrick and Hall 2004). However, in *Drosophila,* the number of cell divisions in the male and female germlines has been shown to be similar (Drost and Lee 1995), suggesting that there should be no significant difference in the mutation rate between the sexes. Nevertheless, other factors may differentially affect X-linked and autosomal mutation rates such as selection on codon usage on the X given a higher level of codon usage bias and GC content on the X across many *Drosophila* species (Singh *et al*. 2005a; b, 2008; Vicoso *et al*. 2008; Campos *et al*. 2013; Schrider *et al*. 2013; Keightley *et al*. 2014). Past studies on *D. melanogaster* have not shown a statistically significant difference between X-linked and autosomal mutation rates (Keightley *et al*. 2009, 2014; Schrider *et al*. 2013). For other species, evidence of a lower X-linked mutation rate is inconclusive, with few species showing evidence of lower X mutation rates (Garrigan *et al*. 2014), however more studies are needed to better understand X-linked versus autosomal mutation rate differences.

Nonetheless, we performed simulations to test the impact of a lower mutation rate on the X and whether this could, on its own, explain the patterns observed in the data. Simulations show that only the most extreme case of lower X-linked mutation rate (*µ_X_* =0.1*µ_A_*) can decrease diversity below the 20% of π_X,Neutral_, where π_X,Neutral_ is the average π/bp from neutral X chromosome simulations. However, none of our low X-linked mutation rate simulations significantly elevated haplotype homozygosity. In fact, the only case that can considerably reduce π_X,_ *µ_X_* =0.1*µ_A_,* shows the lowest haplotype homozygosity across all models (**Fig. 4C**). This is due to longer windows in terms of base pairs for the same SNP window size used across all scenarios which leads to a lower probability of sampling two identical haplotypes and hence a lower haplotype homozygosity. From the above, we conclude that it is unlikely that the low diversity windows on the X are uniquely explained by a lower X-linked mutation rate.

### Demography

Next, we tested the effects of demography on X versus autosome diversity, as population bottlenecks can have different effects on the X versus autosomes due to differences in *N_e_*. We tested two variations of a bottleneck model: a severe and short bottleneck and a shallow and long bottleneck (**Methods**). Both models were fit to π/bp and S/bp in short introns from the Drosophila Genetic Reference panel (DGRP) dataset (Garud *et al*. 2015, 2021). Short introns, known to evolve almost neutrally, were defined as introns shorter than 86 bp with the first 16 bp and last 6 bp removed (Clemente and Vogl 2012; Lawrie *et al*. 2013). Our simulations revealed a π_X_/π_A_ slightly below 0.75 as well as a significant increase in haplotype homozygosity in both bottleneck scenarios (**Fig. 4A,B**). To test whether the higher variance on the X, expected from its smaller *N_e_,* could result in local strong dips in diversity we looked at whether the distribution of π_X_ could achieve values below 20% of the average π_X_ for each model. In **Fig. 4D** we see that the 20% of the average π_X_ is well below the tail of the π_X_ distribution, suggesting that bottlenecks are unlikely to generate the strong local dips in π that we observe in the data (**Fig. 2**).

Finally, inbreeding is another process that could result in local depletion of diversity. However, if this were the case, we would expect to see similar reductions in diversity across the autosomes, which we do not observe. Nonetheless, removal of closely related individuals in the *D. melanogaster* populations (**Methods**) still results in depleted nucleotide diversity on the X relative to autosomes.

### The effect of background selection on nucleotide diversity and haplotype homozygosity

Having investigated neutral evolutionary scenarios, we next examined whether selection can generate the patterns observed in the data. Both background selection (BGS) and selective sweeps can decrease genetic diversity at linked sites, either through the purging of neutral alleles that are linked to deleterious mutations or through hitchhiking, in which a beneficial mutation and its genetic background spread rapidly through the population (Tajima 1989; Charlesworth *et al*. 1993, 1995; Stephan 2010). However, background selection is not known to increase haplotype homozygosity (Enard *et al*. 2014; Schrider 2020).

To test whether BGS can, on its own, give rise to values of π_X_/π_A_<0.75, decrease π/bp below the low diversity threshold and elevate haplotype homozygosity, we simulated a population (*Ne=*1x10^6^) in which a fraction (*d=*0.1,0.5 and 0.8) of mutations are deleterious. The selection coefficient (*s_d_*) for the deleterious mutations followed a gamma distributed DFE with mean and shape parameter -0.000133 and 0.35, respectively (Huber *et al*. 2017). Our simulations show that (i) BGS does not decrease π_X_ such that π_X_/π_A_ is considerably below 0.75 (**Fig. 5A**) (ii) diversity is reduced below 20% of π_X,Neutral_ only when 80% of mutations are deleterious (**Fig. 5B**) and (iii) none of the BGS models considered can elevate homozygosity (**Fig. 5C**).

**Figure 5.**
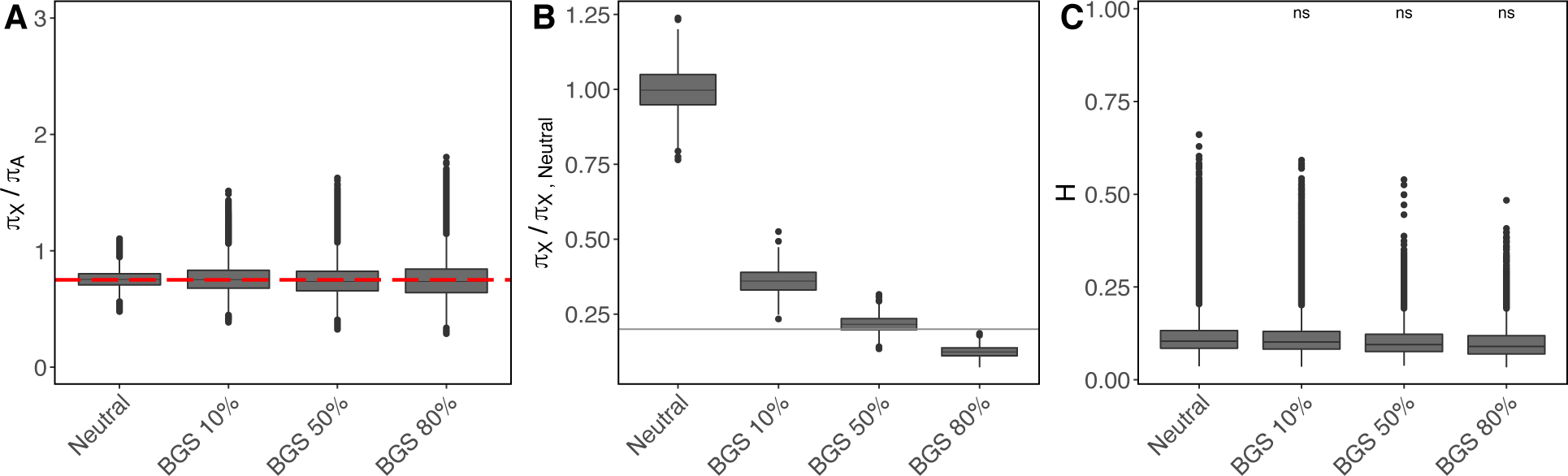
Effect of background selection on diversity and haplotype homozygosity. The models considered include: a neutral scenario with no sex ratio or mutation rate biases and three BGS models. For the BGS models, we varied the proportion of deleterious mutations (10, 50 and 80%) while the DFE for the deleterious selection coefficient (*s_d_*) was gamma distributed with mean and shape parameter -0.000133 and 0.35, respectively (Huber *et al*. 2017). For each model, we computed: **(A)** π_X_/π_A_, where the red dashed line corresponds to the expected π_X_/π_A_ = 0.75 value in a completely neutral case; **(B)** π_X_/ π_X,Neutral_ where π_X,Neutral_ is the π in the constant *N_e_* neutral model with no biases and the solid gray line is 20% of π_X,Neutral_; and **(C)** haplotype homozygosity. The asterisks represent a significant elevation in haplotype homozygosity relative to neutrality using a one-sided Wilcoxon rank-sum test.

### The combined effect of background selection, demography, X-linked mutation rates and recombination rate variation on nucleotide diversity and haplotype homozygosity

Separately, each of the variables we have examined thus far (BGS, reduced *µ_X_*, low recombination rates, and bottlenecks) cannot produce strong dips in diversity and elevated haplotype homozygosity. However, in combination, these variables may be able to generate the patterns observed in the data. To test this, we simulated the full combination of variables, including BGS, bottlenecks, reduced *µ_X_* and low recombination rates, which individually showed some, but not all, of the signatures observed in the data. We did not simulate sex bias because female bias elevates diversity on the X (opposite trend of what is observed in 5 of 7 populations) and male bias is not reported to be a dominant process influencing the populations under study.

Figure 6 shows the result of combining BGS, bottlenecks and low *µ_X_* for *r* = 5e-7cM/bp, while **Figure S8** shows these results for a lower recombination rate (*r*=2.5e-7cM/bp). We found that when *r* = 5e-7cM/bp and 50% or more of new mutations are deleterious, only one of the bottlenecks displayed dips in diversity below the designated low diversity threshold. However, none of these scenarios showed elevated haplotype homozygosity. By contrast, when *r* = 2.5e- 7cM/bp, a single scenario was able to recapitulate all the signatures from the data: a shallow long bottleneck with 50% of deleterious mutations and *µ_X_/µ_A_*=0.9. However, empirical evidence of lower X-linked mutation rates in *Drosophila* is inconclusive (see section *Mutation rates* above). Moreover, we note that species with sample sizes of a similar order of magnitude as those of our simulations (*D. melanogaster* and *D. simulans*) exhibit a more pronounced elevation in haplotype homozygosity than what we observe in **Figure S8**. This indicates a larger effect size in our data compared to simulations, where due to the large number of replicates we can detect smaller effect sizes as statistically significant.

**Figure 6.**
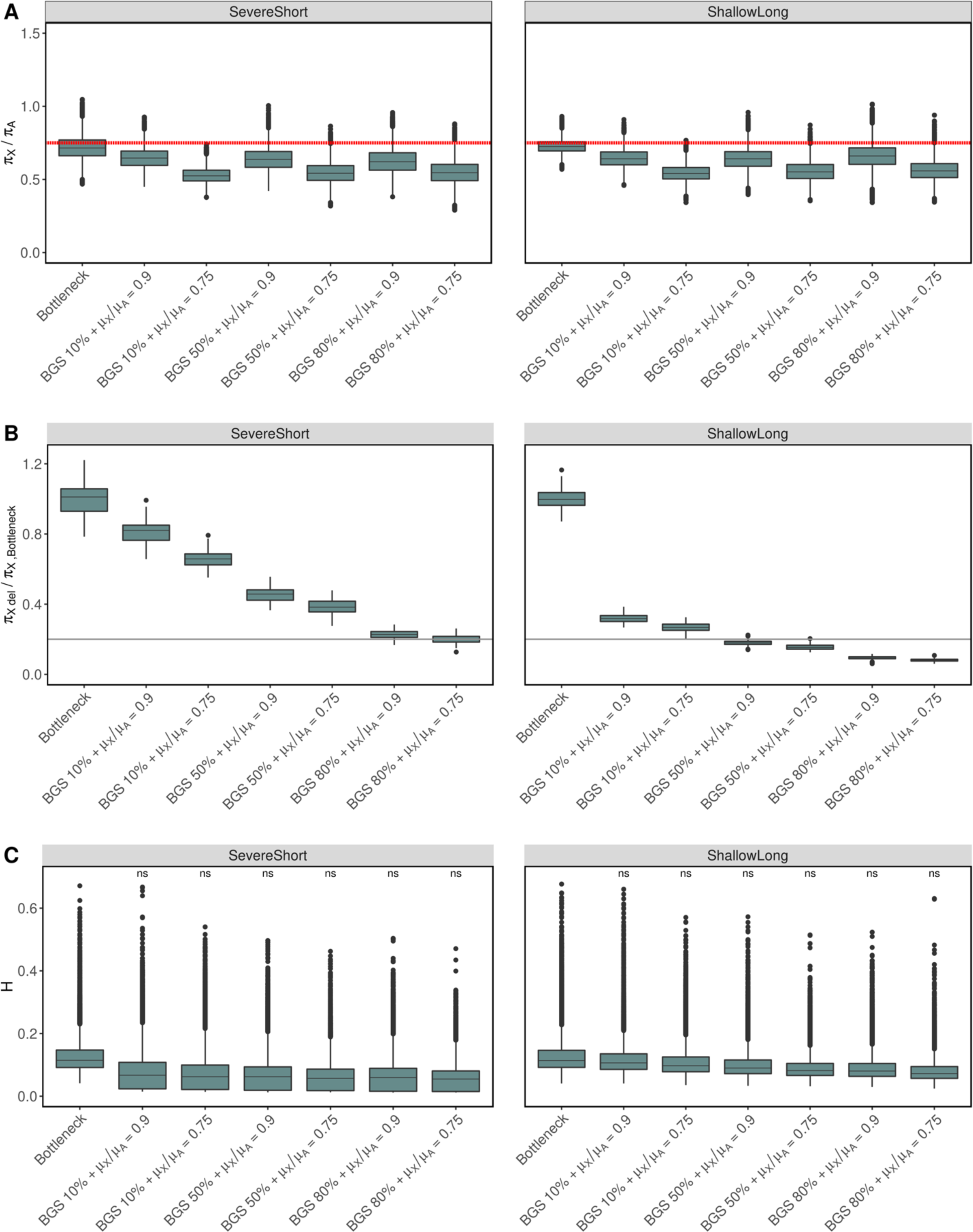
Effect of background selection combined with bottlenecks and low X-linked mutation rate on diversity patterns. The models considered include the combination of a bottleneck model with one of three BGS models each combined with a lower X linked mutation rate (µ_X_*/*µ_A_ =0.9, µ_X_*/*µ_A_=0.75). For the BGS models, we varied the proportion of deleterious mutations (10, 50 and 80%) while the DFE for the deleterious selection coefficient (*s_d_*) was gamma distributed with mean and shape parameter -0.000133 and 0.35, respectively (Huber *et al*. 2017). The recombination rate was set to *r*=5e-7 cM/bp across all simulations. Figure S8 shows the same corresponding plot for half the recombination rate (2.5e-7 cM/bp). The left column corresponds to a severe short bottleneck model and the right column to a shallow long bottleneck model (**Methods**). For each model, we computed: **(A)** π_X_/π_A_ across all models, where the red dashed line corresponds to the expected π_X_/π_A_ = 0.75 value in the neutral model, **(B)** π_X_/π_X_,_Bottleneck_ where π_X_,_Bottleneck_ is the average π in the corresponding bottleneck model with no added biases and the solid gray line is 20% of π_X_,_Bottleneck_, **(C)** Haplotype homozygosity across all models and results from a one-sided Wilcoxon rank-sum test for elevation of H in the bottleneck+BGS+Low µ_X_ scenario compared to a neutral bottleneck model.

Finally, to test whether the effect of background selection is stronger when a larger number of deleterious mutations reside on the same chromosome, we simulated longer chromosomes of 1Mb instead of 20Kb (**Fig. S9**). These simulations included BGS for three different recombination rates (**Methods**). We found that when recombination is low (*r*<=1e-8 cM/bp), diversity dips below 20% of π_X,Neutral_ and haplotype homozygosity is elevated. In this scenario, recombination is not effectively breaking linkage, leading to an elevation of haplotype homozygosity and strong reductions in diversity compared to π_X,Neutral._ However, in this scenario, π_X_/π_A_ does not dip below 0.75, which is inconsistent with five of seven populations in the data (**Fig. S9**). Additionally, we note that the fraction of the low diversity windows in the *D. melanogaster* genome that experiences a recombination rate below *r*=1e-8cM/bp is 0.15, making this unlikely to be the driving force generating dips in diversity.

### Hard sweeps can generate the patterns observed on the X across species

Our results indicate that the patterns observed in the data are unlikely to be generated by sex bias, low recombination rates, low X-linked mutations rates, demography or BGS, either individually or in combination. Next, we tested the effect of positive selection in generating dips in diversity and elevated haplotype homozygosity on the X chromosome. To do so, we simulated hard and soft sweeps (**Methods**), varied the strength of selection (*N_e_s_b_ =* 20, 200 and 2000) and computed π_X_/π_A_, π_X_/π_X,Neutral_ and haplotype homozygosity in a 20Kb window for each scenario.

Our simulations show that selective sweeps can decrease π_X_/π_A_ below 0.75 and elevate haplotype homozygosity (**Fig. 7A, C**), but only hard sweeps can reduce diversity below 20% of π_X,Neutral_ (**Fig. 7B**) as long as selection is sufficiently strong (*Nes_b_* =2000). Moreover, compared to the scenarios that we have simulated thus far, we observe a much stronger elevation in haplotype homozygosity when there is positive selection. This is indicative of a more substantial effect size, which aligns better with observations in *D. melanogaster* and *D. simulans,* the species with sample sizes comparable to those from simulations.

**Figure 7.**
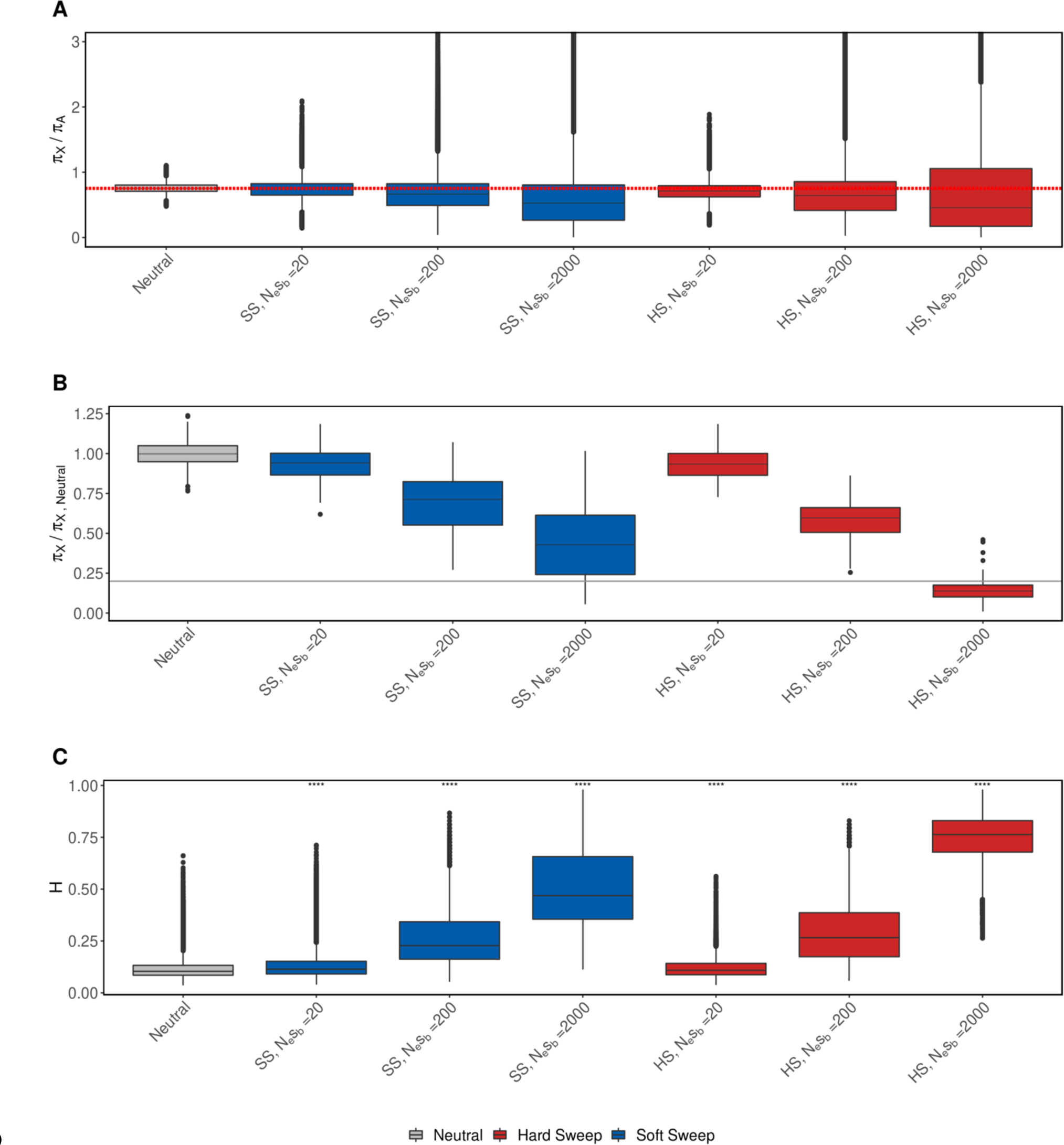
Effect of hard and soft selective sweeps on diversity and haplotype homozygosity. The models considered include: a neutral scenario with no sex ratio or mutation rate biases, three soft sweep (SS) and three hard sweep models (HS). We simulated soft sweep (blue) and hard sweep (red) models (Methods) varying the selection strength of the adaptive mutation (*N_e_s_b_=*20, 200, 2000). For each model, we computed: **(A)** π_X_/π_A_, where the red dashed line corresponds to the expected πX/πA = 0.75 value in a completely neutral case; **(B)** π_X_/ π_X,Neutral_ where π_X,Neutral_ is the π in the baseline neutral model and the solid gray line is 20% of π_X,Neutral_; and **(C)** haplotype homozygosity. The asterisks represent a significant elevation in haplotype homozygosity relative to complete neutrality using a one-sided Wilcoxon rank-sum test.

We note however, that it is also possible for a strong soft sweep that is not too soft (*N_e_s_b_ =2000,* θ_a_ = 0.1 corresponding to approximately two sweeping haplotypes **Fig. S10A**) to reach levels of diversity below the low diversity threshold. However, in this regime (θ_a_ = 0.1) only slightly more than half (∼62%) of the sweeps simulated with θ_a_ = 0.1 are soft (**Fig. S10**) and, among those that are soft, 48% of the simulations show π_X_ below the low diversity threshold compared to 82% for the hard sweep model (**Fig. S11**), making hard sweeps a more likely explanation of the data. Thus, we conclude that patterns of reduced diversity and elevated haplotype homozygosity on the X chromosome of the species analyzed are most consistent with hard sweeps.

## Discussion

A classic question in evolutionary biology is how the evolution of the X chromosome differs from that of autosomes given the X chromosome’s central role in speciation, brain function, fertility, and sexual dimorphism (Rice 1984; Saifi and Chandra 1999; Skuse 2005; Dean and Mank 2014; Payseur *et al*. 2018). Past work has suggested that the X chromosome may exhibit different evolutionary dynamics from the autosomes due to its unique inheritance pattern and increased exposure to selection through male hemizygosity (Vicoso and Charlesworth 2006; Nam *et al*. 2015; Charlesworth *et al*. 2018; Muralidhar and Veller 2022; Harris and Garud 2023). Recently, we found evidence in a North American *D. melanogaster* population that the X chromosome experiences an enrichment of hard selective sweeps compared to autosomes due to the increased visibility of new deleterious mutations to natural selection on the hemizygous X of males and reduced effective population size on the X (Harris and Garud 2023). However, it is unclear whether this pattern of enrichment of hard sweeps on the X is a universal feature across all heterogametic species.

To understand if the enrichment of hard sweeps on the X is common across species, we analyzed multiple whole genome sequences from six *Drosophila* species and found evidence that suggests that hard sweeps are in fact more common on the X chromosome than autosomes across these species. Additionally, we found that the observed patterns of diversity are inconsistent with background selection, differences in mutation rate, population bottlenecks, or sex bias. We note, however, that when combined, in one specific scenario, low recombination rates, low mutation rates, and BGS could generate both strong dips in diversity and elevated haplotype homozygosity (**Fig. S8**). However, in this scenario we observed a small effect size for the elevation of haplotype homozygosity inconsistent with the data (**Fig. 3**, **Fig. S8**). Additionally, we note that there has not been any definitive evidence of reduced mutation rates on the X relative to autosomes in *Drosophila*.

Furthermore, we found that soft sweeps generally cannot generate the patterns observed in the data, except for when soft sweeps are not too soft (e.g. there are only 2 sweeping haplotypes). However, in this scenario (θ_a_ =0.1), only 62% of the simulations led to soft sweeps, and of those, only 48% generated dips in diversity falling below the low diversity threshold, whereas 82% of hard sweeps can. Therefore, we acknowledge that it is possible that other factors other than hard sweeps could be responsible for generating some of the low diversity windows observed in the data. However, our results suggest that these forces are unlikely to be the dominant processes driving the patterns in the data. Future work will be important for disentangling the effects of these forcers on diversity on the X.

The finding that hard sweeps are a likely explanation for the patterns observed on the X chromosome aligns with recent theoretical work predicting harder sweeps on the X chromosome (Muralidhar and Veller 2022; Harris and Garud 2023), as well as empirical work in apes (Nam *et al*. 2015) and *D. mauritiana* (Garrigan *et al*. 2014). In our analysis, we employed a wide range of statistics, including single-site (e.g. π/bp) and multi-site (e.g. haplotype homozygosity) statistics that are sensitive to signatures of selection and also leverage the additional resolution that whole genome sequences provide over genotype data. Combining single site and multi-locus statistics is still a relatively novel area of work that has the potential to reveal patterns of evolution that cannot be detected by either type of statistic alone (Lin *et al*. 2011; Schrider and Kern 2016; Sheehan and Song 2016; Ragsdale and Gutenkunst 2017; Johri *et al*. 2020; Garud *et al*. 2021). Our approach allows us to disentangle the effects of positive selection from other forces, such as background selection and demographic processes, and rule out inconsistent models.

Comparative population genetics, in which multiple genomes from several species each are considered simultaneously, is also a relatively new area of work given the paucity of deep population genetic samples from multiple species. Population genetic studies have traditionally focused on analyzing multiple genomes from a single species, e.g. (Arbiza *et al*. 2014; Garud and Rosenberg 2015; Signor *et al*. 2018) with few examining more than two population genetic whole genome datasets from different species simultaneously ((Nam *et al*. 2015; Chen *et al*. 2018; Nadachowska-Brzyska *et al*. 2019; Latrille *et al*. 2023; Rodrigues *et al*. 2023) with (Latrille *et al*. 2023) restricted to exomes), leaving open numerous avenues of inquiry on the commonalities and idiosyncrasies of population genetic processes.

Now, with the increasing availability of deep population genetic sequences from multiple species, we can examine population genetic processes across many species. In this study, the ability to compare multiple species reveals that the extent of positive selection on the X chromosome may not be equal across all species. For example, we observed few low diversity windows on the X in *D. teissieri*, while in *D. santomea* we observed a low diversity region extending up to ∼200Kb (**Fig. 2**). Additionally, this comparative approach reveals deviations from the trend in individual species. One species that in particular looked very different was *D. sechellia*. This species showed a unique trend in which π_X_/π_A_ was the highest in exons and lowest in intergenic regions, even after stringent filtering of the data (**Methods**; **Fig. 1**). A potential explanation for this behavior could be abundant introgression from *D. simulans* to *D. sechellia*, as previously reported (Garrigan *et al*. 2012; Matute and Ayroles 2014; Schrider *et al*. 2018). Interestingly, introgression appears to be less common on the X chromosome, potentially due to the involvement of sex chromosomes in hybrid incompatibilities (Maroja *et al*. 2015; Turissini and Matute 2017; Schrider *et al*. 2018). Moreover, functional regions of the genome are less likely to exhibit evidence of introgression, with exons being the least susceptible and intergenic regions having higher rates of introgressed regions (Sankararaman *et al*. 2014). As a result, if introgression is abundant, the difference in diversity between the X chromosome and autosomes should be more pronounced in intergenic regions and least in exons, potentially driving the observed trend in the data.

Despite individual species showing deviations from the trend, some species exhibited commonalities. For example, the species analyzed in this study all had a higher proportion of low diversity regions on the X chromosome compared to the autosomes. Additionally, we found some targets of selection to be shared across species, however they were generally few in number (**Fig. S12-S14**, **Text S1**). Instead, most genes found in low diversity windows were found in a single species only, indicating that while hard sweeps might be common on the X, different functions may be under selection in different species. We note that this finding is distinct from that of primates (Nam *et al*. 2015) where there appears to be more overlap across species. This suggests that the underlying mechanisms of selection in *Drosophila* versus primates may differ.

Our paper is a clear demonstration of how comparative population genetic analyses can reveal new insights into the forces that shape genetic variation within and across species, which we expect to serve as a useful example as new high-resolution population genomic datasets from multiple species become increasingly available in the coming years. While our study did not examine the prevalence of soft sweeps in autosomes and the X chromosome due to limited samples sizes per species, future research with larger sample sizes could provide the relevant data needed to be able to detect soft sweeps with haplotype homozygosity statistics, thereby providing a more comprehensive understanding of the tempo and mode of adaptation across species (Pennings and Hermisson 2006b). Ultimately, our study highlights the significance of hard sweeps in shaping the diversity patterns of the X chromosome across species and suggests an important evolutionary mechanism that may be widespread among all species. Future work may reveal the potential role of these hard sweeps in driving sexual dimorphism and speciation given the X chromosome’s significant involvement in these important processes.

## Data availability

Sequence data for *D. melanogaster* can be downloaded from www.johnpool.net. For *D. simulans* the vcf file can be downloaded from https://zenodo.org/record/154261#.YzMzty2z3jC For the remaining species, sequence data are available at NCBI’s Short Read Archive with accession numbers given in Table S1.

## Acknowledgments

We are grateful to Rebekah Rogers and Peter Andolfatto for providing the *D.simulans* w501 genome assembly. We thank Dmitri Petrov and Kirk Lohmueller for helpful conversations. MH was supported by the National Institutes of Health Systems and Integrative Biology Training Grant (NIH-NIGMS 5T32GM008185) and the National Institutes of Health Training Grant in Genomic Analysis and Interpretation (NIH T32HG002536). BYK was supported by NIH grant NIGMS F32GM135998. NRG was supported by funding from a National Science Foundation CAREER award.

## Supplemental text

**Table S1.**
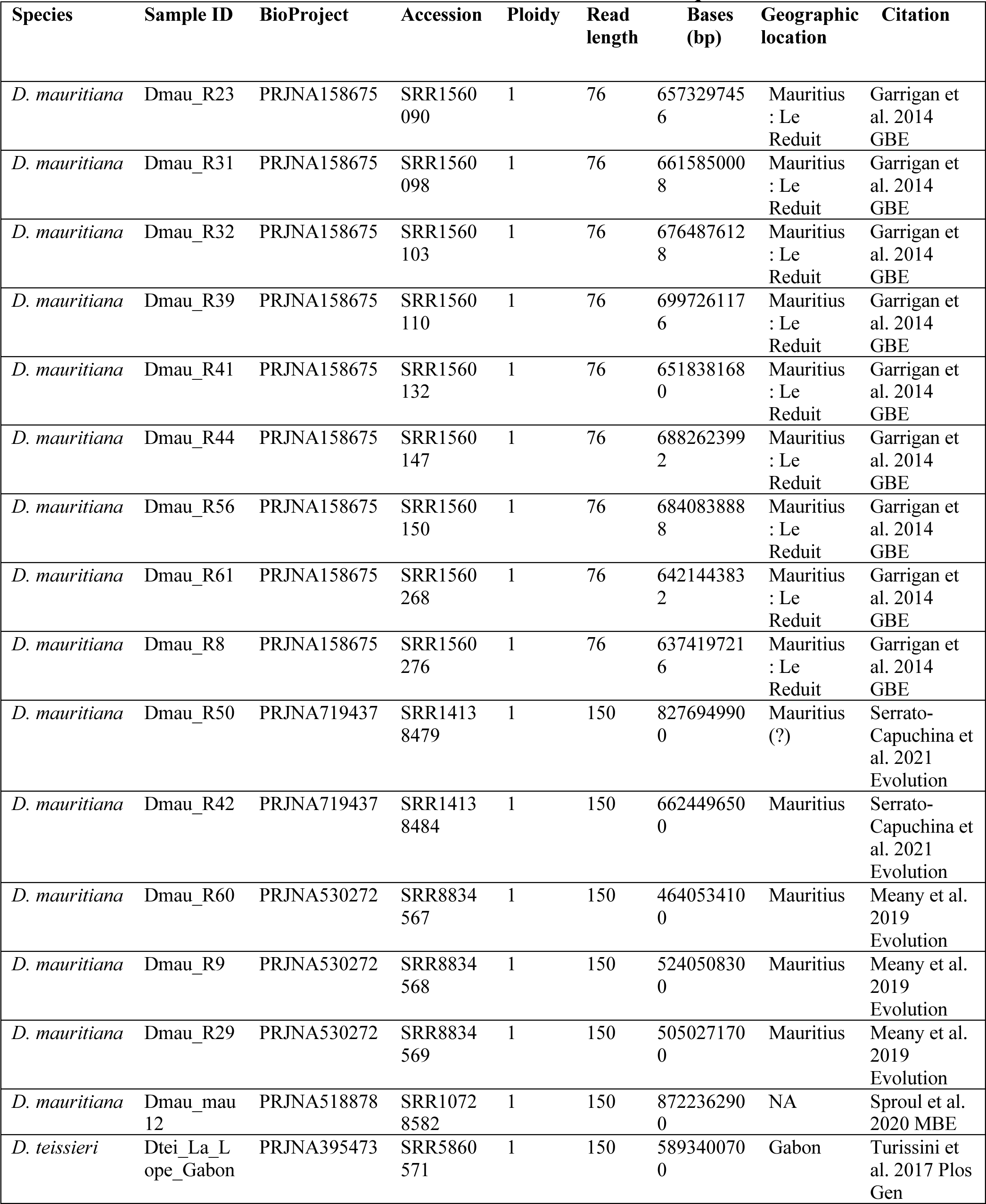
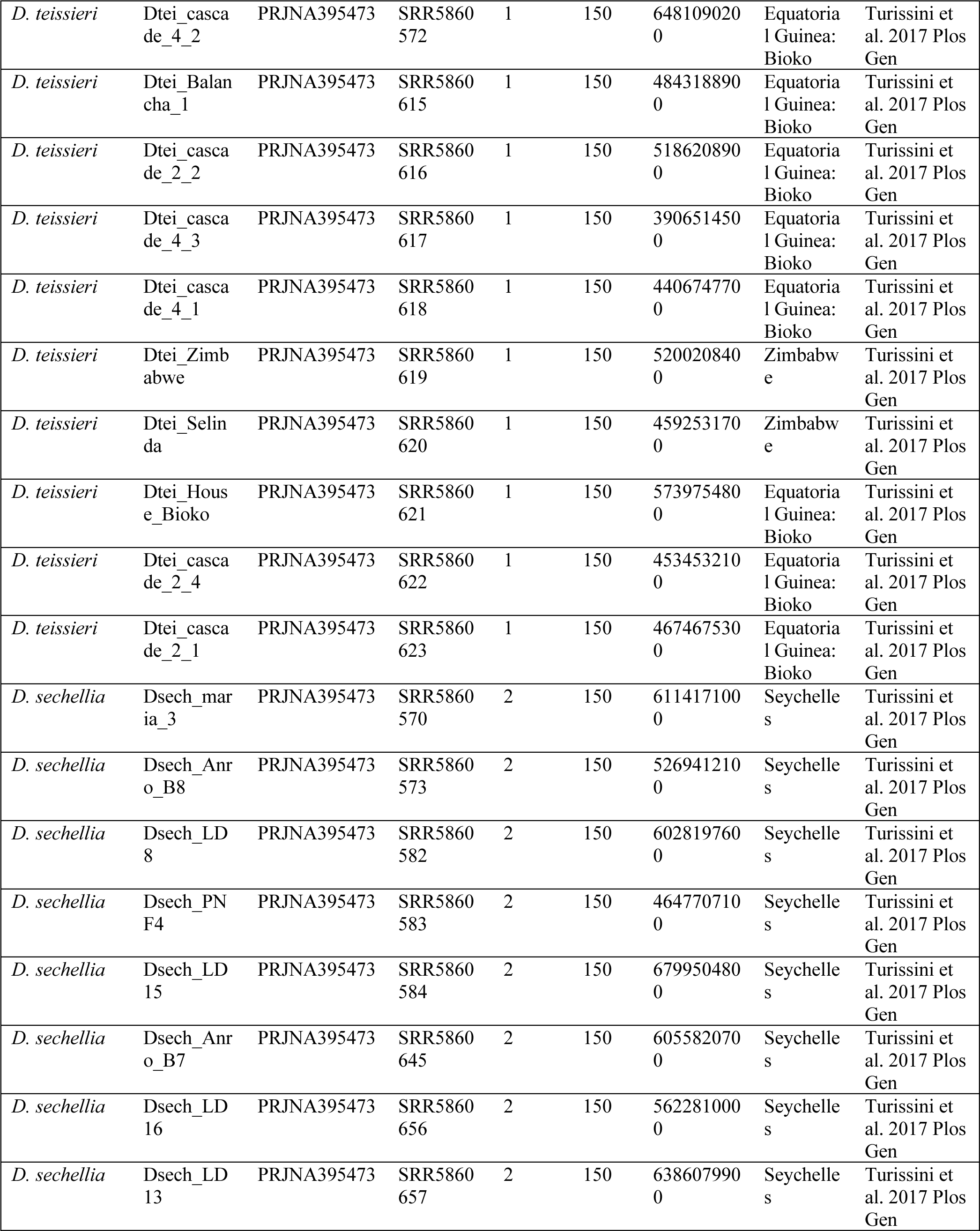
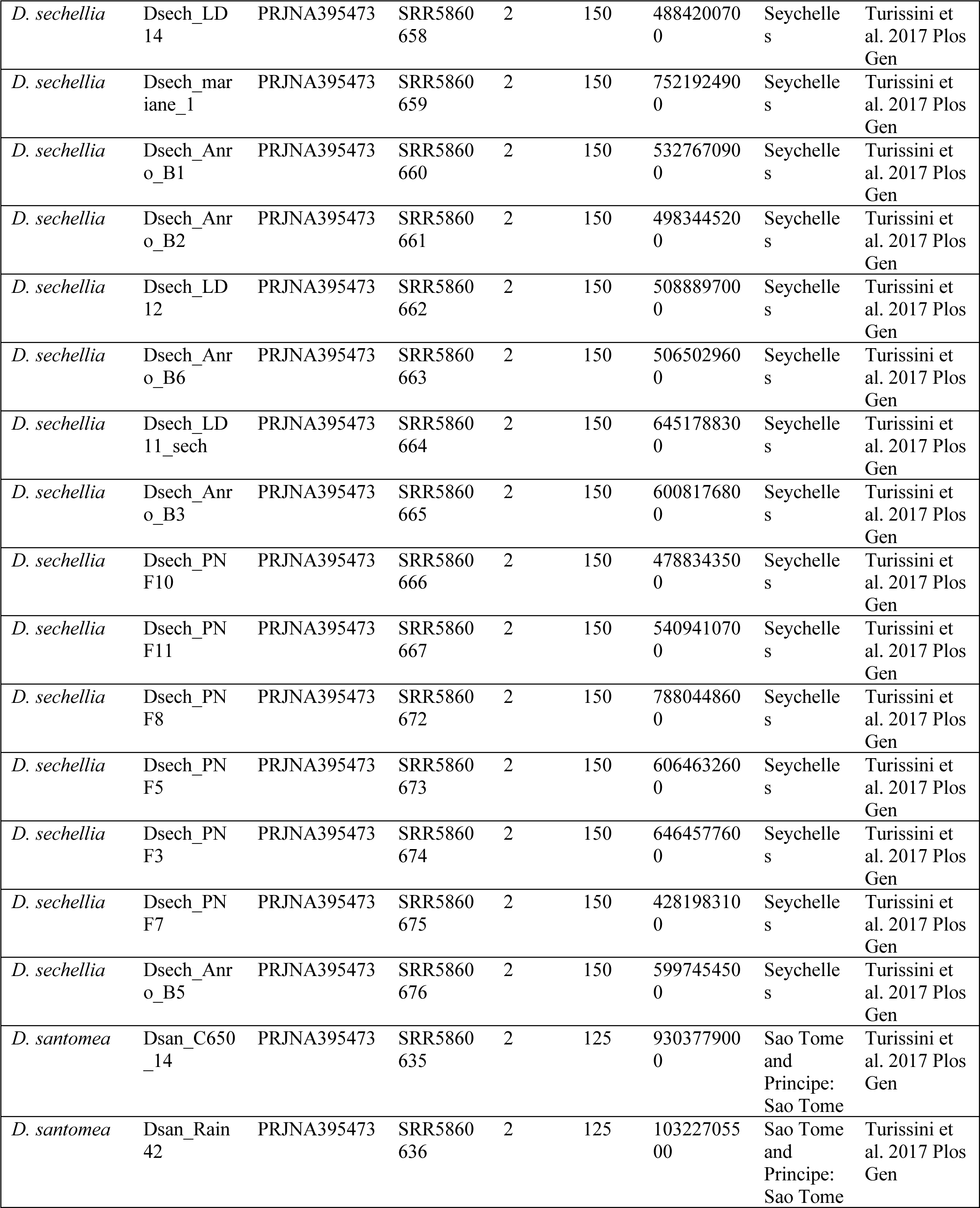
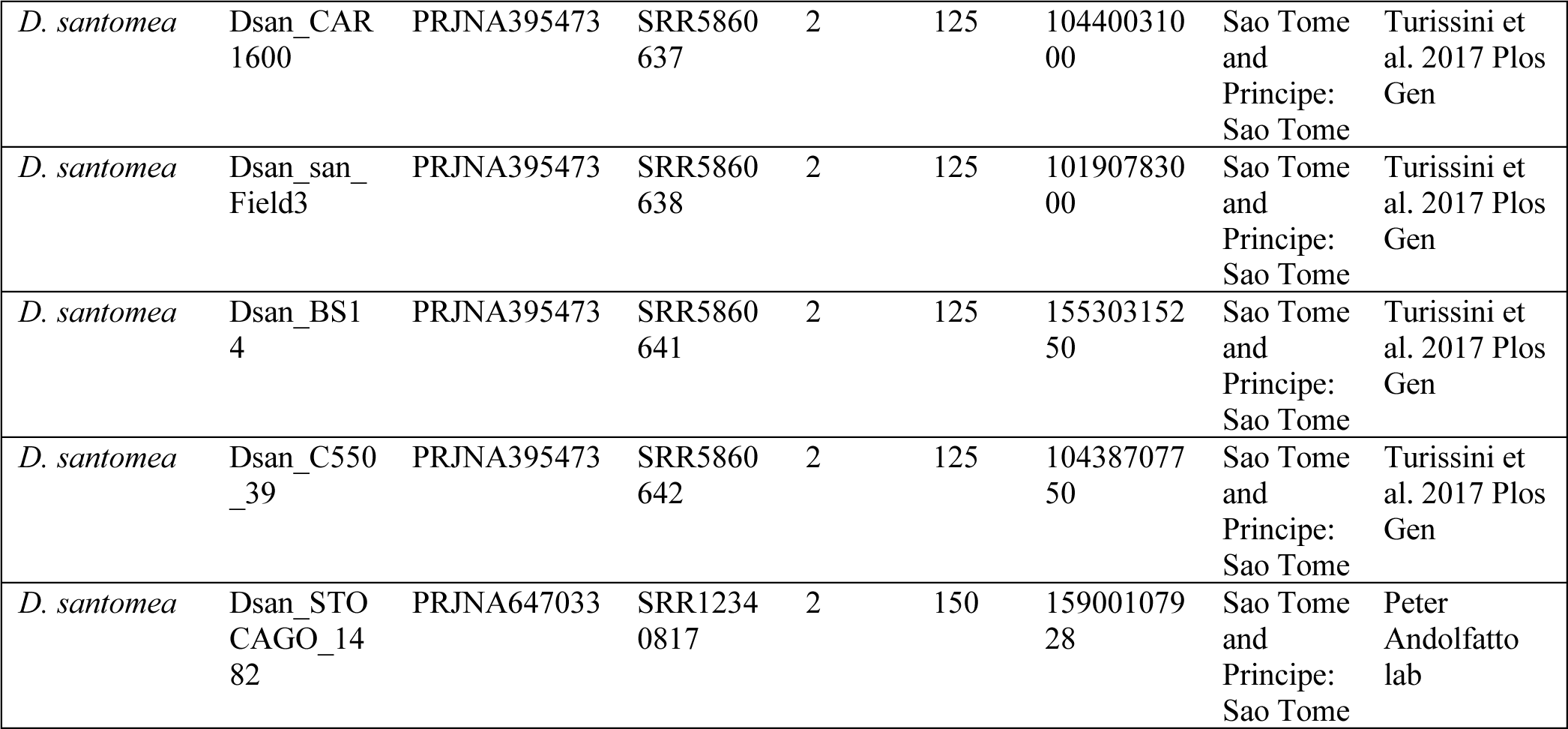
D.mauritiana, D. teissieri, D. sechellia and D. santomea sequences.

**Table S2.** Drosophila species and populations used in study. Clade, population of origin, number of samples, and reference are given in each column.

| **Species** | **Clade** | **Population** | **# Samples used** | **Source** |
| --- | --- | --- | --- | --- |
| *D. melanogaster* | *melanogaster* | North America (RAL) | 100 | MacKay et al. 2012;<br>Lack et al 2015 |
| *D. melanogaster* | *melanogaster* | Africa (ZI) | 100 | Pool et al 2012;<br>Lack et al 2015 |
| *D. simulans* | *simulans* | North America (CA) | 170 | Signor 2018 |
| *D. sechellia* | *simulans* | Africa (Praslin) | 23 | Schrider et al. 2018 |
| *D. mauritiana* | *simulans* | Africa (Mauritius) | 15 | Garrigan et al. 2014<br>Serrato-Capuchina et al.<br>2021<br>Meany et al. 2019 |
| *D. santomea* | *yakuba* | Africa (São Tomé) | 7 | Turissini and Matute<br>2017;<br>Bachtrog et al 2007; |
| *D. teissieri* | *yakuba* | Africa (Bioko, Zimbabwe,<br>Gabon) | 11 | Turissini and Matute<br>2017 |

**Table S3.** Minimum and mean π/bp computed in non-overlapping 20Kb windows on the X chromosome across species. The fourth column shows the minimum π/bp on the X as a percent of the average π/bp on the X chromosome. In *D. simulans* the window with lowest diversity is 0.5% of the average π/bp while on *D. santomea* it is 10% of the average π/bp.

| Species | min($\pi$/bp) | mean($\pi$/bp) | % of mean($\pi$/bp) |
| --- | --- | --- | --- |
| *D.mauritiana* | 0.00004 | 0.006 | 0.7% |
| *D. melanogaster (RA)* | 0.0002 | 0.003 | 6.7% |
| *D. melanogaster (ZI)* | 0.0006 | 0.008 | 7.5% |
| *D.santomea* | 0.0005 | 0.005 | 10.0% |
| *D. sechellia* | 0.000002 | 0.0003 | 0.7% |
| *D. simulans* | 0.00002 | 0.004 | 0.5% |
| *D. teissieri* | 0.0003 | 0.01 | 3.0% |

**Figure S1.**
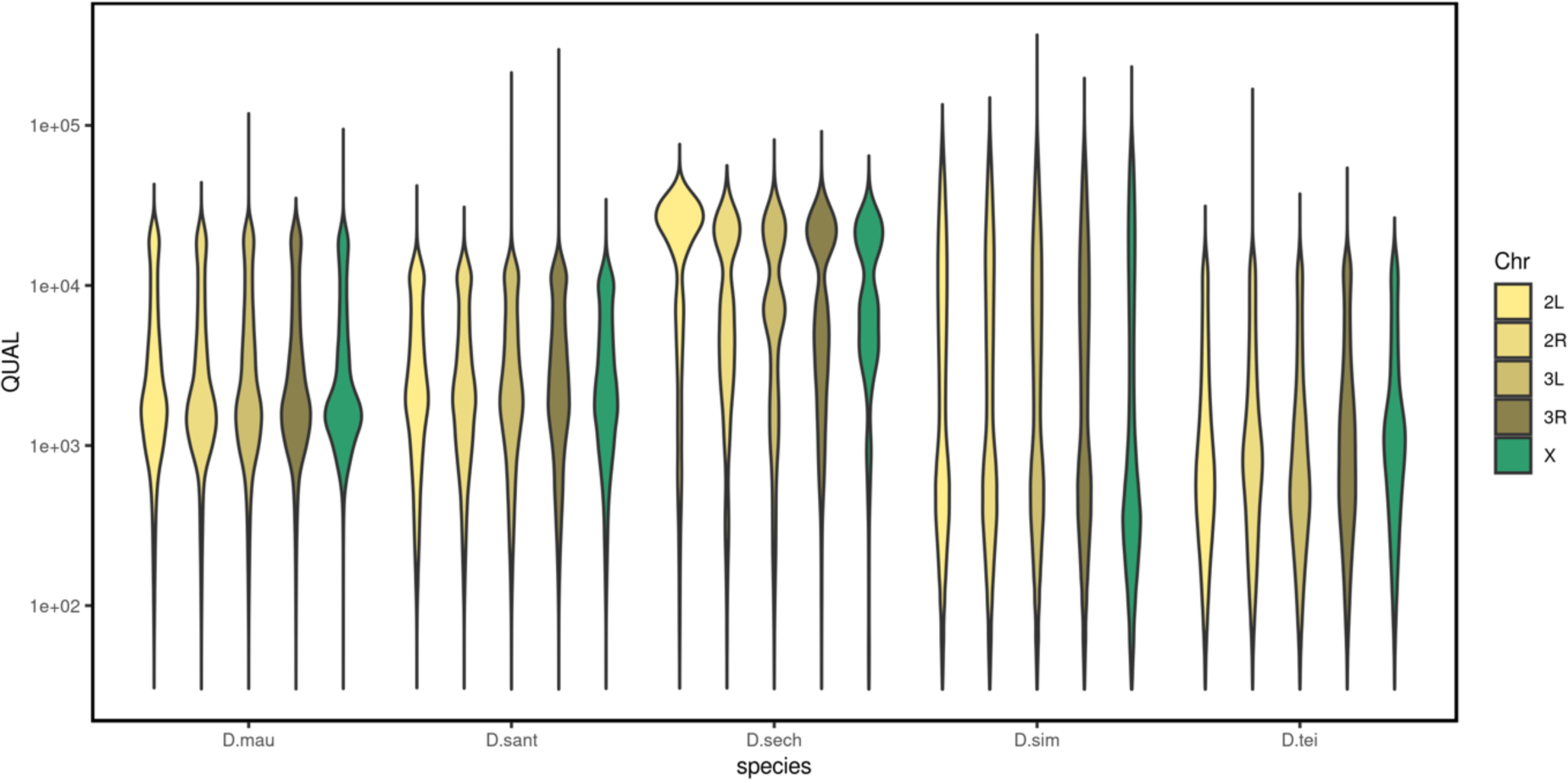
Phred-scaled quality score across species. The distribution is similar across autosomes and the X chromosome for *D. mauritiana, D. santomea. D. sechellia, D. simulans* and *D. teissieri*. The y-axis is log-scaled for better visualization.

**Figure S2.**
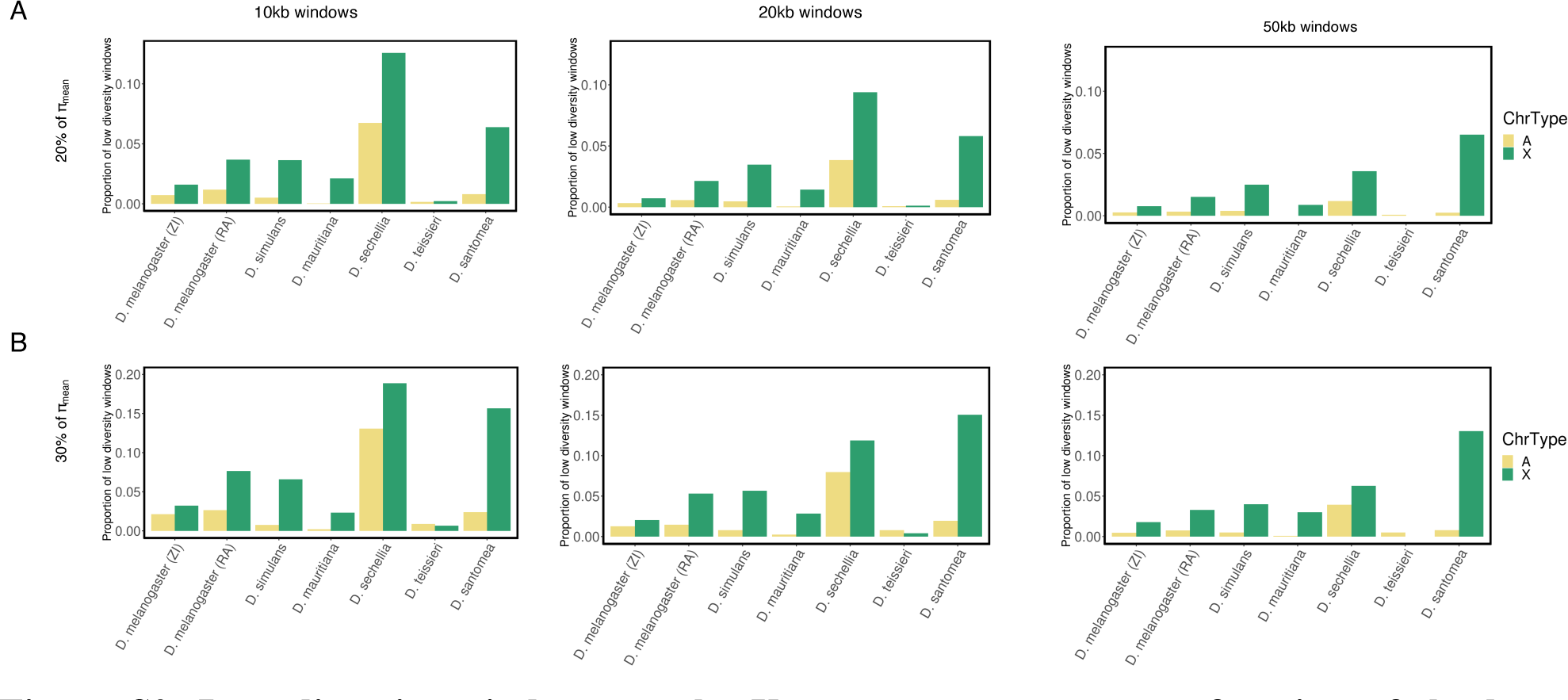
Low diversity windows on the X versus autosomes as a function of the low diversity thresholds and window size. Proportion of low diversity windows on X versus autosomes (top row, green and yellow bars). Three different analysis windows were tested: 10kb, 20kb and 30kb windows, as well as two different diversity thresholds: **(A)** 20% and **(B)** 30% of the chromosome’s π/bp average. We consistently see a higher proportion of low diversity windows on the X compared to autosomes. This figure is analogous to **Fig. 2B** in the main text.

**Figure S3.**
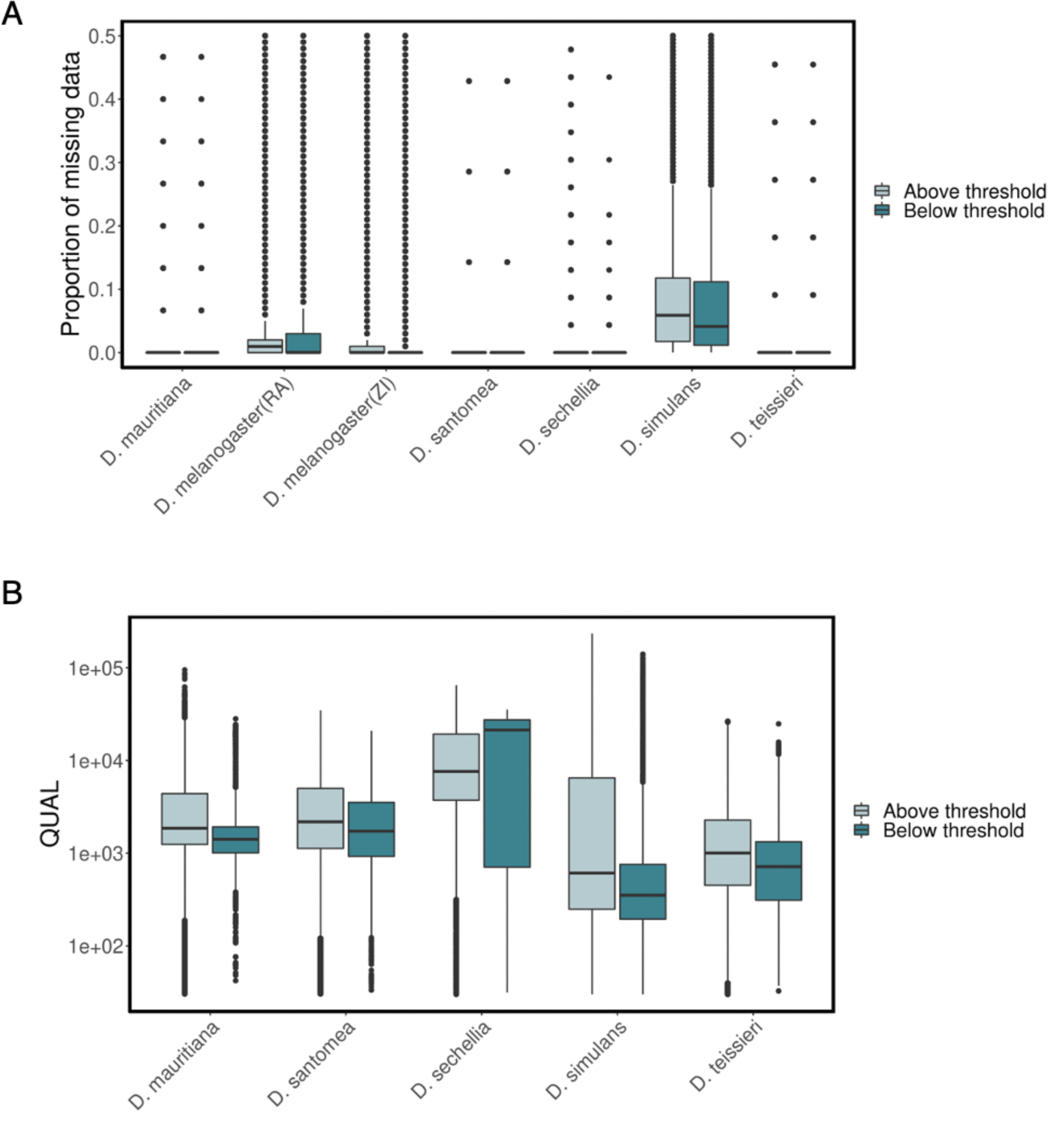
Proportion of missing data (A) and Phred-scaled quality score (B) in windows above and below the low diversity threshold for the X chromosome. The y-axis in (B) is log-scaled for better visualization.

**Figure S4.**
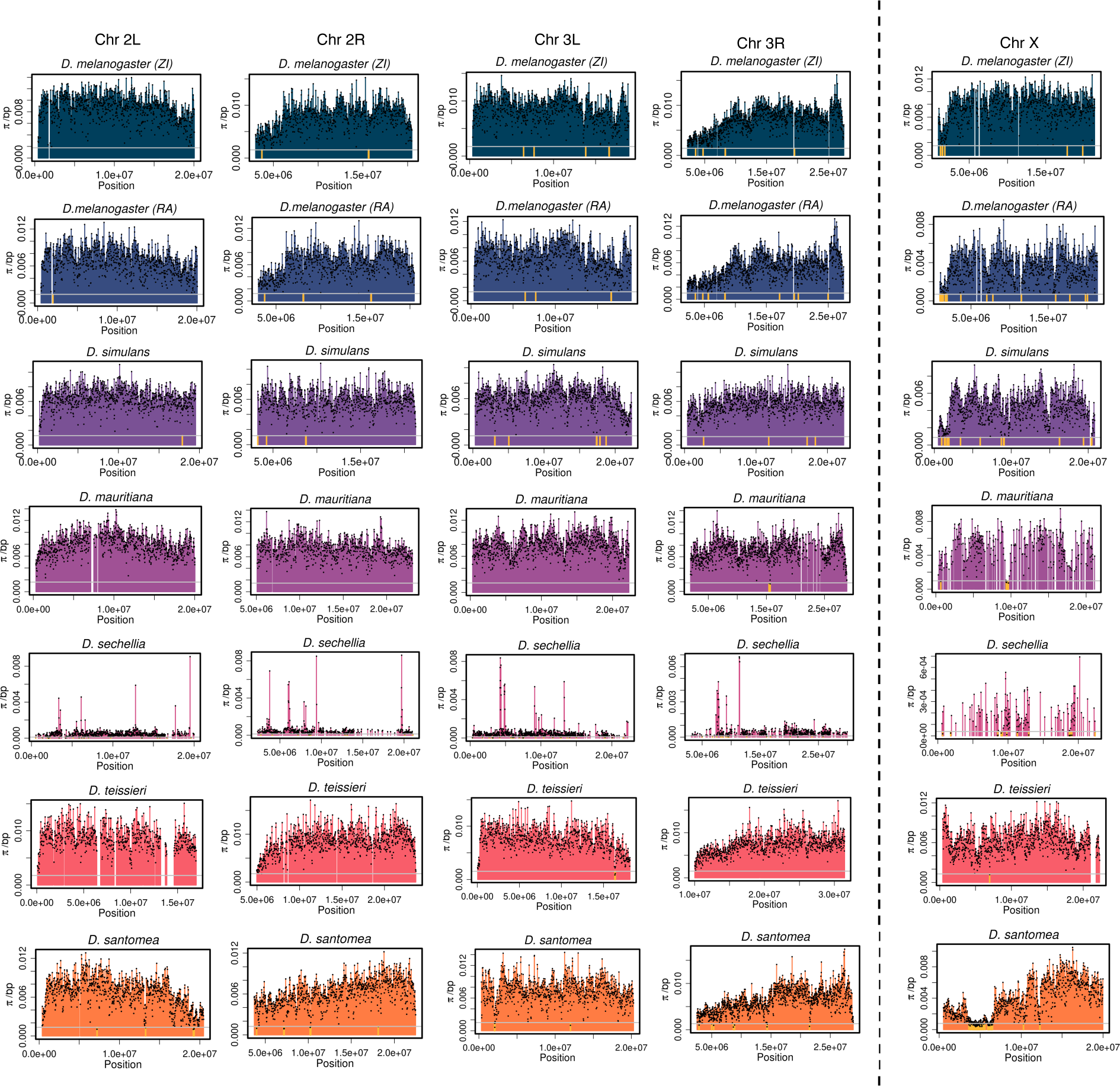
Genome-wide diversity in six species. π/bp in 20kb bins across four autosomal arms and the X chromosome of six *Drosophila* species and two *D. melanogaster* populations from Raleigh (RA) and Zambia (ZI). Windows where diversity is below 20% of the chromosomal average (grey horizontal line) are highlighted in gold.

**Figure S5.**
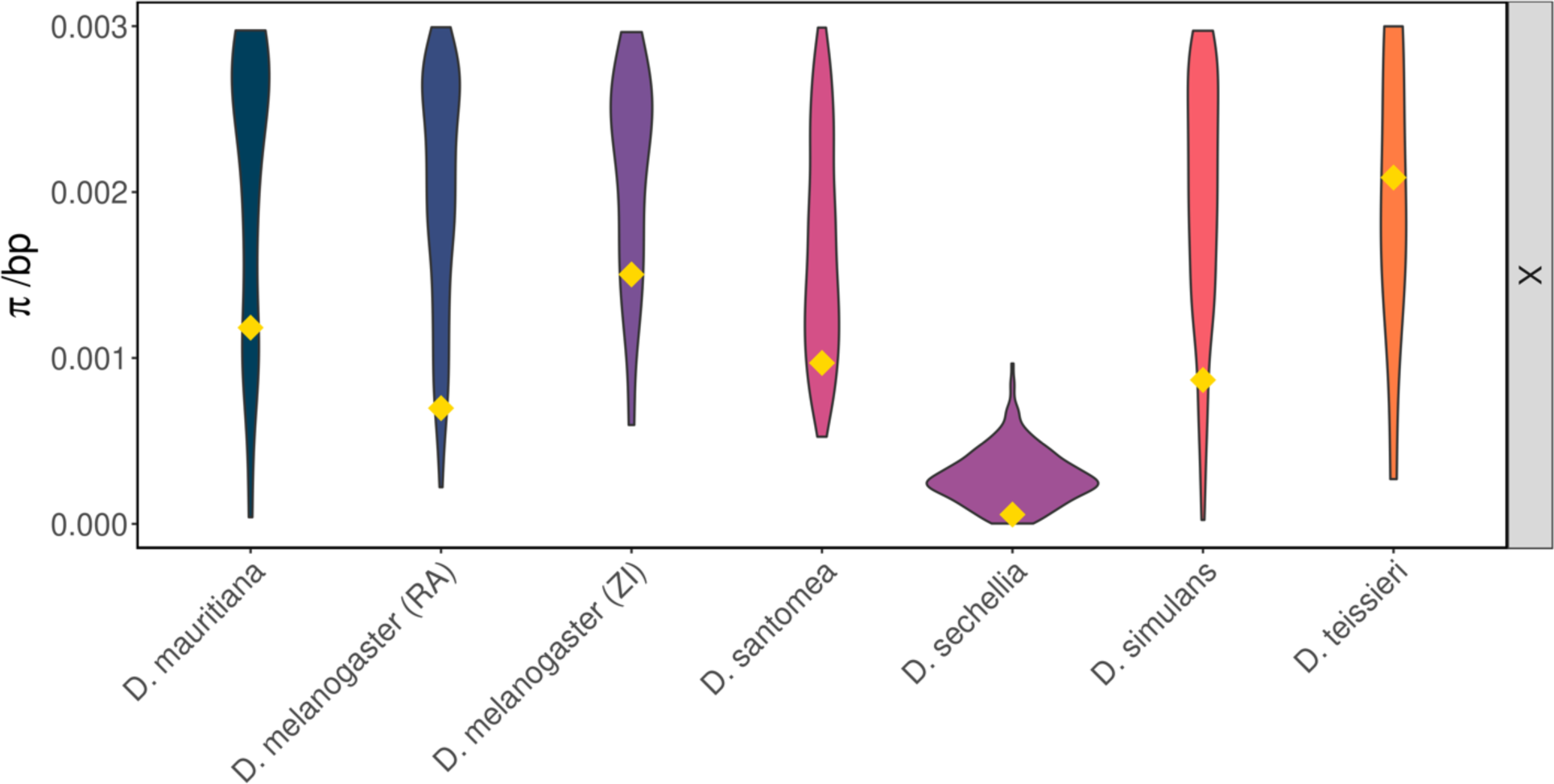
Distribution of π/bp for the X chromosome across species. Violin plots for π/bp for each species’ X chromosome are plotted with y-axis going up to π/bp=0.03 for better visibility of the lower tail of the distribution. The low diversity threshold is shown as the yellow diamond on the tail of each violin plot and represents 20% of the mean value of the respective distribution. The lower tail of each distribution extends well below the low diversity threshold for all species.

**Figure S6.**
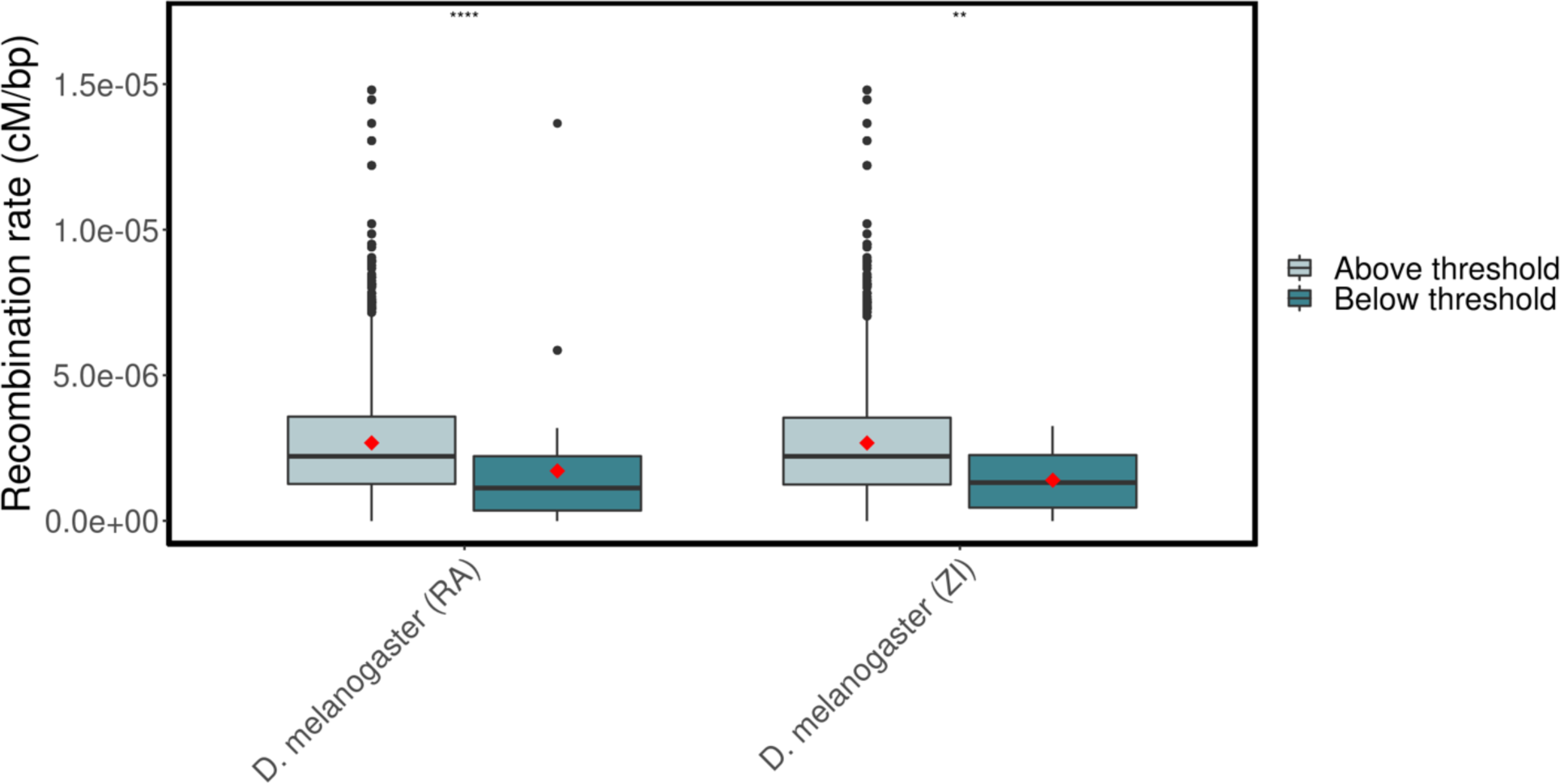
*D. melanogaster* recombination rates (cM/bp) in regions below and above the low diversity threshold. Recombination rates for *D. melanogaster* were obtained from the Comeron et al 2012 recombination map. The red data points correspond to the average recombination rate in each category (r = 1.71 x 10-6 cM/bp (RA),1.40 x 10-6 cM/bp (ZI) in low diversity windows and r = 2.68 x 10-6 cM/bp (RA), 2.67 x 10-6 cM/bp (ZI) in high diversity windows). We find a lower recombination in low diversity windows (dark blue) compared to windows above (light blue) the low diversity threshold. The asterisks represent a significant difference in recombination rate using a Wilcoxon rank-sum test.

**Figure S7.**
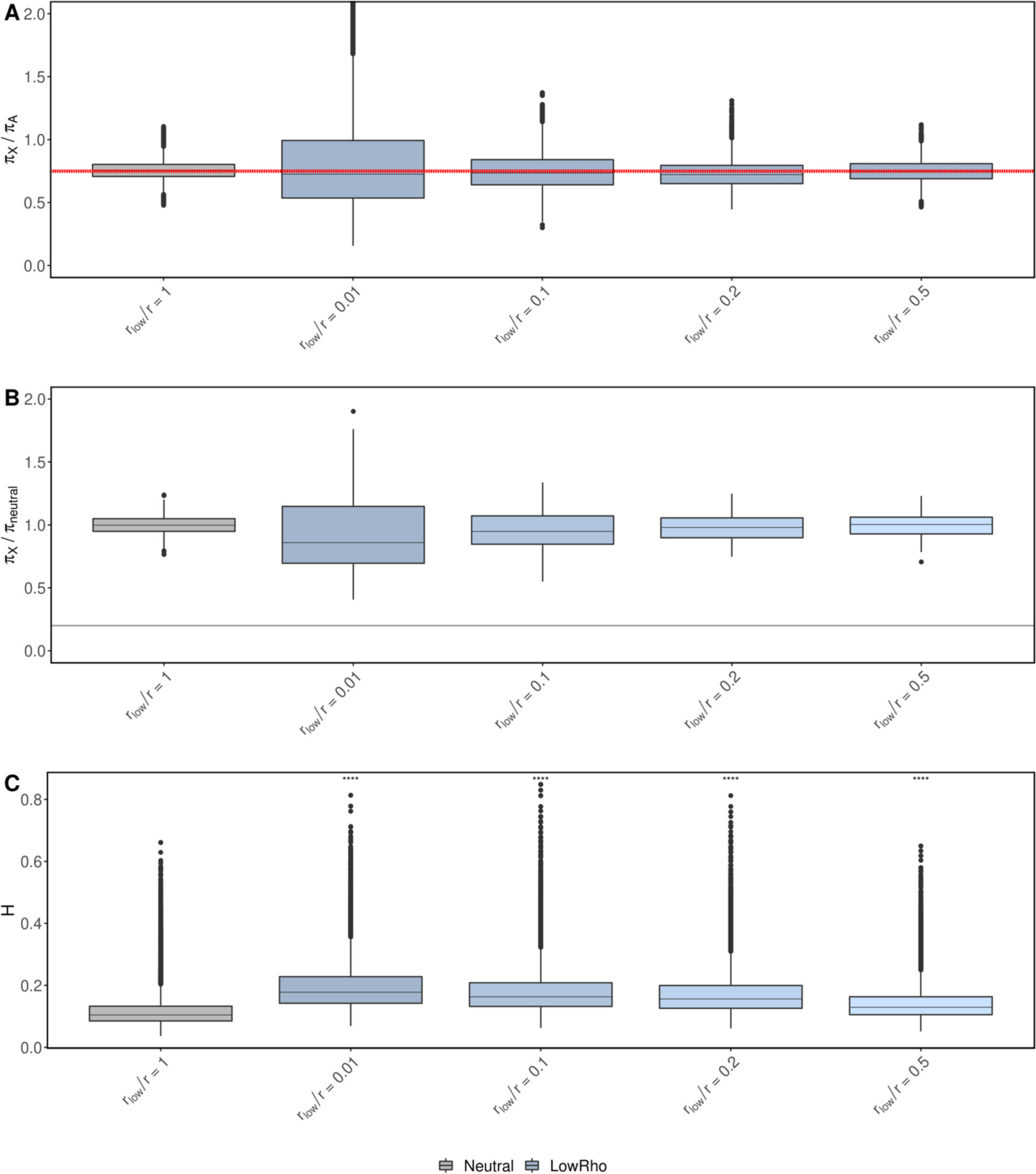
Windows of low recombination can elevate haplotype homozygosity but cannot reduce diversity below 20% of π_neutral_. Models of regions with low recombination rate (*r_low_*) such that *r_low_/r=0.01,0.1,0.2* and *0.5,* where *r=5e-9cM/bp.* These regions of low recombination cannot reduce X-linked diversity below the π_X_/π_A_ =0.75 expectation (**A**) or below the low diversity threshold **(B)** but can elevate haplotype homozygosity compared to neutrality **(C).**

**Figure S8.**
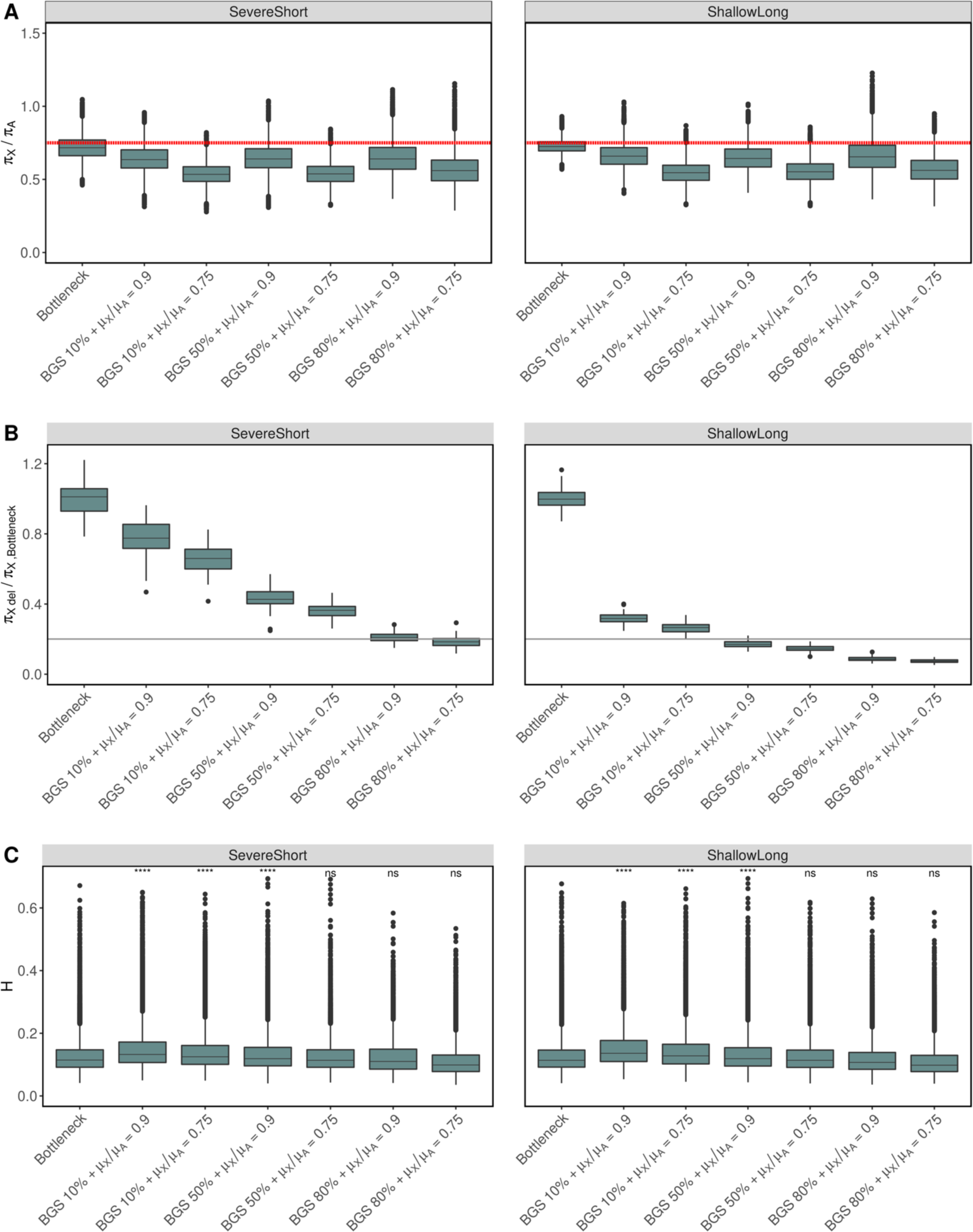
Background selection combined with bottlenecks and low X-linked mutation rate for *r=* 2.5e-7cM/bp. The models considered models considered include three BGS models each combined with a lower X linked mutation rate (µ_X_*/*µ_A_ =0.9, µ_X_*/*µ_A_=0.75). For the BGS models, we varied the proportion of deleterious mutations (10, 50 and 80%) while the DFE for the deleterious selection coefficient (*s_d_*) was gamma distributed with mean and shape parameter - 0.000133 and 0.35, respectively (Huber *et al*. 2017). The left column corresponds to a severe short bottleneck model and the left to a shallow long bottleneck model (**Methods**). For each model, we computed: **(A)** π_X_/π_A_ across all models, where the red dashed line corresponds to the expected π_X_/π_A_ = 0.75 value in the baseline neutral model. **(B)** π_X_/π_X_,_Bottleneck_ where π_X_,_Bottleneck_ is the average π in the corresponding bottleneck model with no added biases. The mean π_X_,_Bottleneck_ represents the average diversity in the chromosome. The solid gray line is 20% of π_X_,_Bottleneck_. **(C)** Haplotype homozygosity across all models and results from a one-sided Wilcoxon rank-sum test for elevation of H in the bottleneck+BGS+Low µ_X_ scenario compared to a neutral bottleneck model.

**Figure S9.**
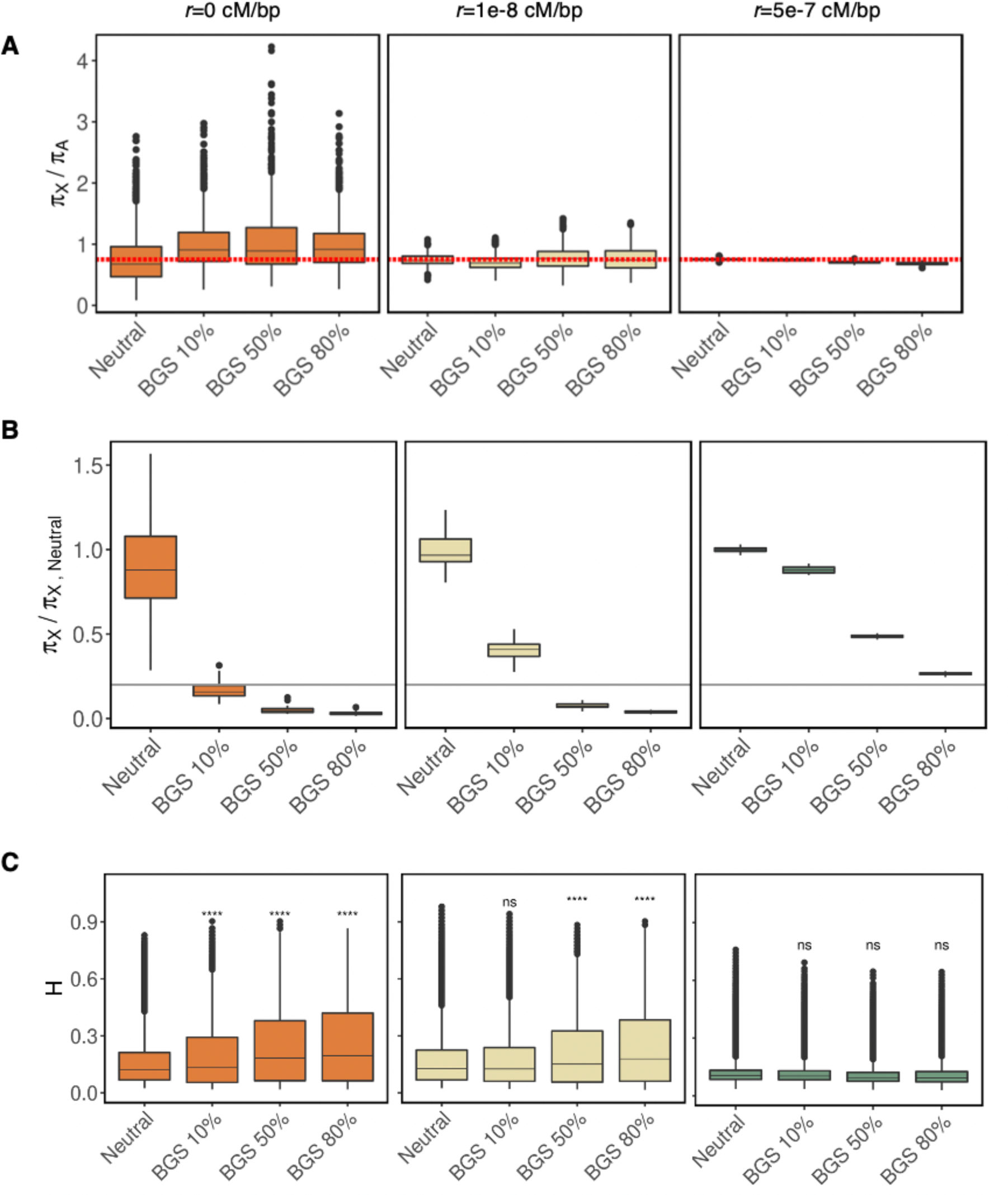
Nucleotide diversity (π) and haplotype homozygosity (H) in 1Mb simulated chromosomes with BGS, varying recombination rate (*r)*. The models considered include a neutral model with a constant *N_e_=10^6^* and no sex bias, and, three BGS models where we varied the proportion of deleterious mutations (10, 50 and 80%). The DFE for the deleterious selection coefficient (s_d_) was gamma distributed with mean and shape parameter -0.000133 and 0.35, respectively. Simulated three different recombination rates: *r*=0,1e-8 and 5e-7 cM/bp. For each model, we computed: **(A)** π_X_/π_A_, where the red dashed line corresponds to the expected πX/πA = 0.75 value in a completely neutral case; **(B)** π_X_/π_X,Neutral_ where π_X_ is the average π in the scenario being considered on the X-axis, and π_X,Neutral_ is the average π in the baseline neutral model. The solid gray line is 20% of π_X,Neutral_; and **(C)** haplotype homozygosity. The asterisks represent a significant elevation in haplotype homozygosity relative to complete neutrality using a one-sided Wilcoxon rank-sum test.

**Figure S10.**
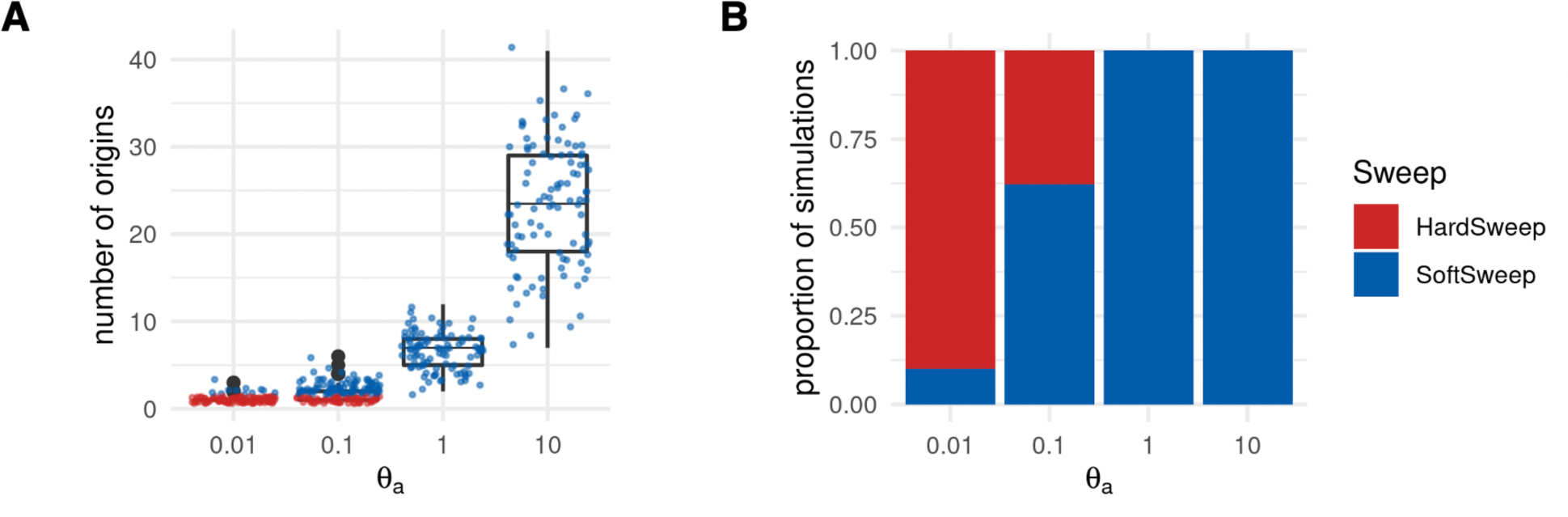
**Number of origins and proportion of hard/soft sweep simulations as a function of θ_a._** Soft sweeps (blue) were defined as simulations in which the sample had >1 mutational origin for the adaptive mutation. Hard sweeps (red) were defined as simulations with one mutational origin for the adaptive mutation. **(A)** Distribution of the number of origins for the adaptive mutation as a function of θ_a._ **(B)** Proportion of hard versus soft sweeps in simulations as a function of θ_a._

**Figure S11.**
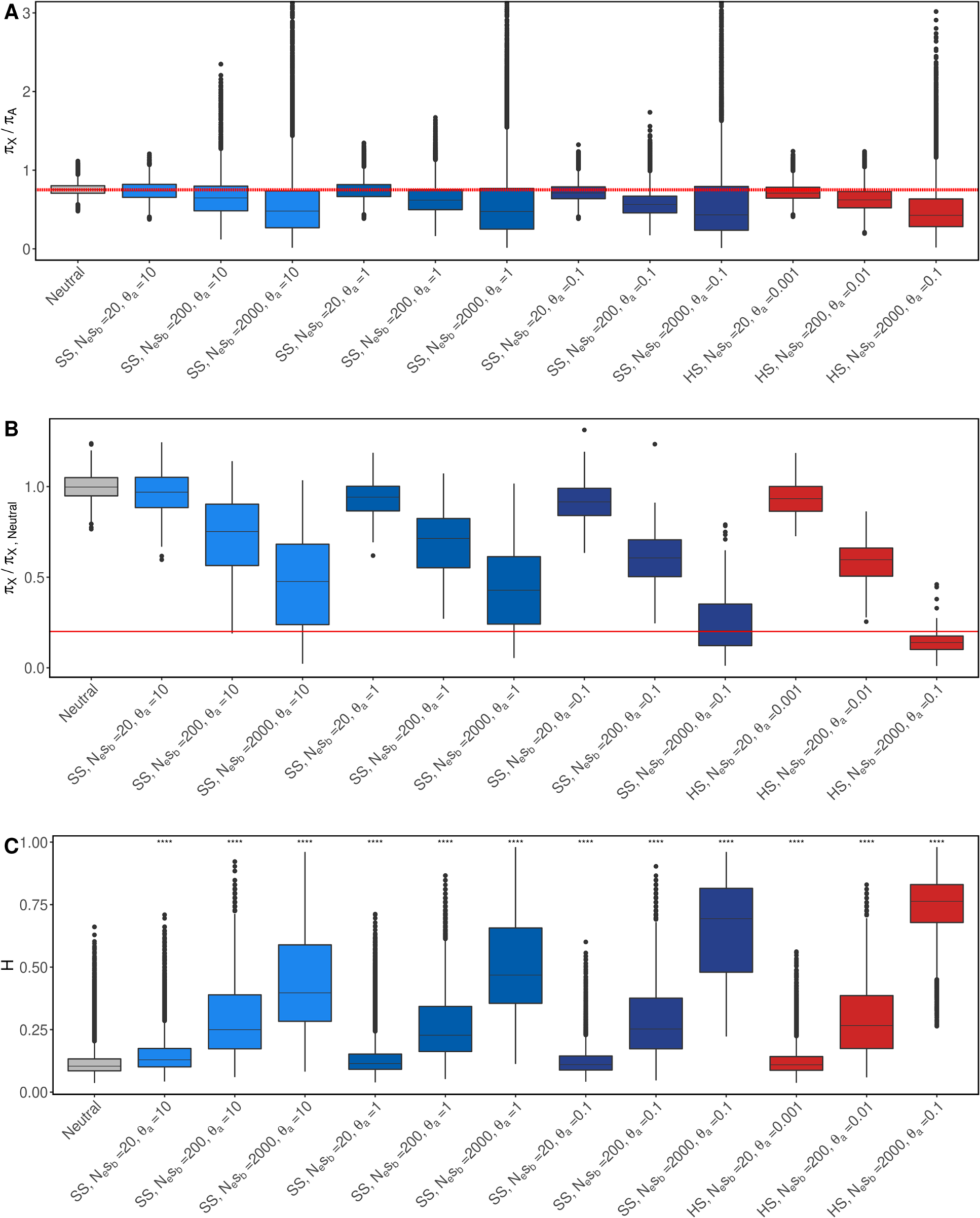
Diversity (π) and haplotype homozygosity (H) in hard and soft sweeps. Models of hard sweeps (red; θ_a_ =0.01) and soft sweeps varying softness (blue; θ_a_ =0.1,1 and 10) with three different selection strengths (*N_e_s_b_=*20, 200, 2000). For each model, we computed: **(A)** π_X_/π_A_, where the red dashed line corresponds to the expected πX/πA = 0.75 value in a completely neutral case; **(B)** π_X_/π_neutral_ where π_neutral_ is the average π in the baseline neutral model and the solid gray line is 20% of π_neutral_; and **(C)** haplotype homozygosity. The asterisks represent a significant elevation in haplotype homozygosity relative to complete neutrality using a one-sided Wilcoxon rank-sum test. Both hard and soft sweeps can decrease π_X_/π_A_ <0.75 and elevate haplotype homozygosity. When selection is strong (*N_e_s_b_=*2000) it is possible for both hard and soft sweeps to reduce π_X_ below 20% of π_neutral_. However, the proportion of hard sweeps simulations below this threshold is higher (0.82) compared to soft sweeps (0.48 for θ_a_ =0.1, 0.19 for θ_a_ =1 and 0.18 for θ_a_ =10), making hard sweeps more likely.

**Figure S12.**
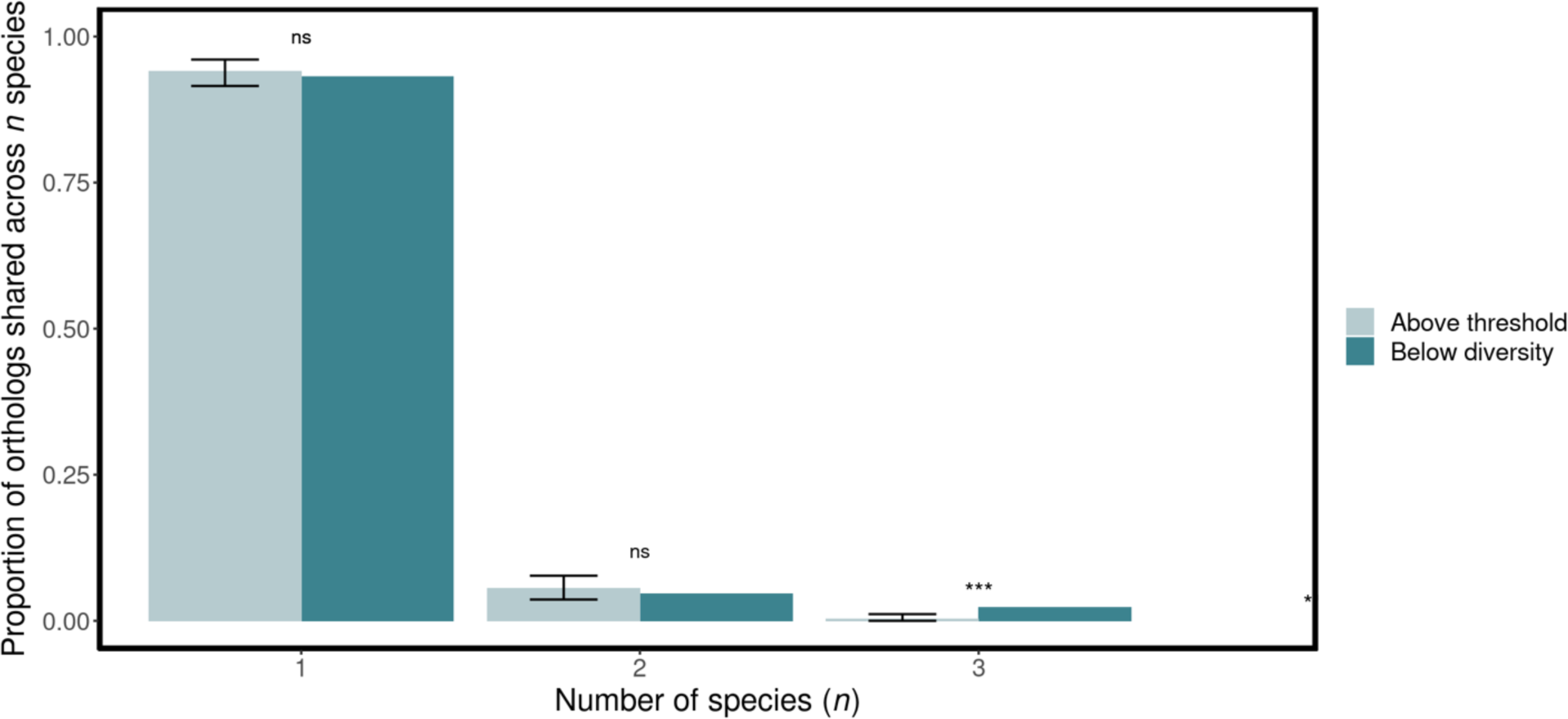
Proportion of orthologs shared across species in windows below and above the low diversity threshold. We obtained 100 random samples of orthologs found in windows above the low diversity threshold and obtained the mean number of orthologs shared across *n=1,2 or 3* species (light blue) as well as the corresponding 95% confident intervals for each case. The case of *n=1* represents orthologs found in a single species only. For *n=3,* we found that the proportion of orthologs shared across species is higher in low diversity windows. Comparisons between the two *D. melanogaster* populations were excluded from the analysis in order to emphasize between-species dynamics.

**Figure S13.**
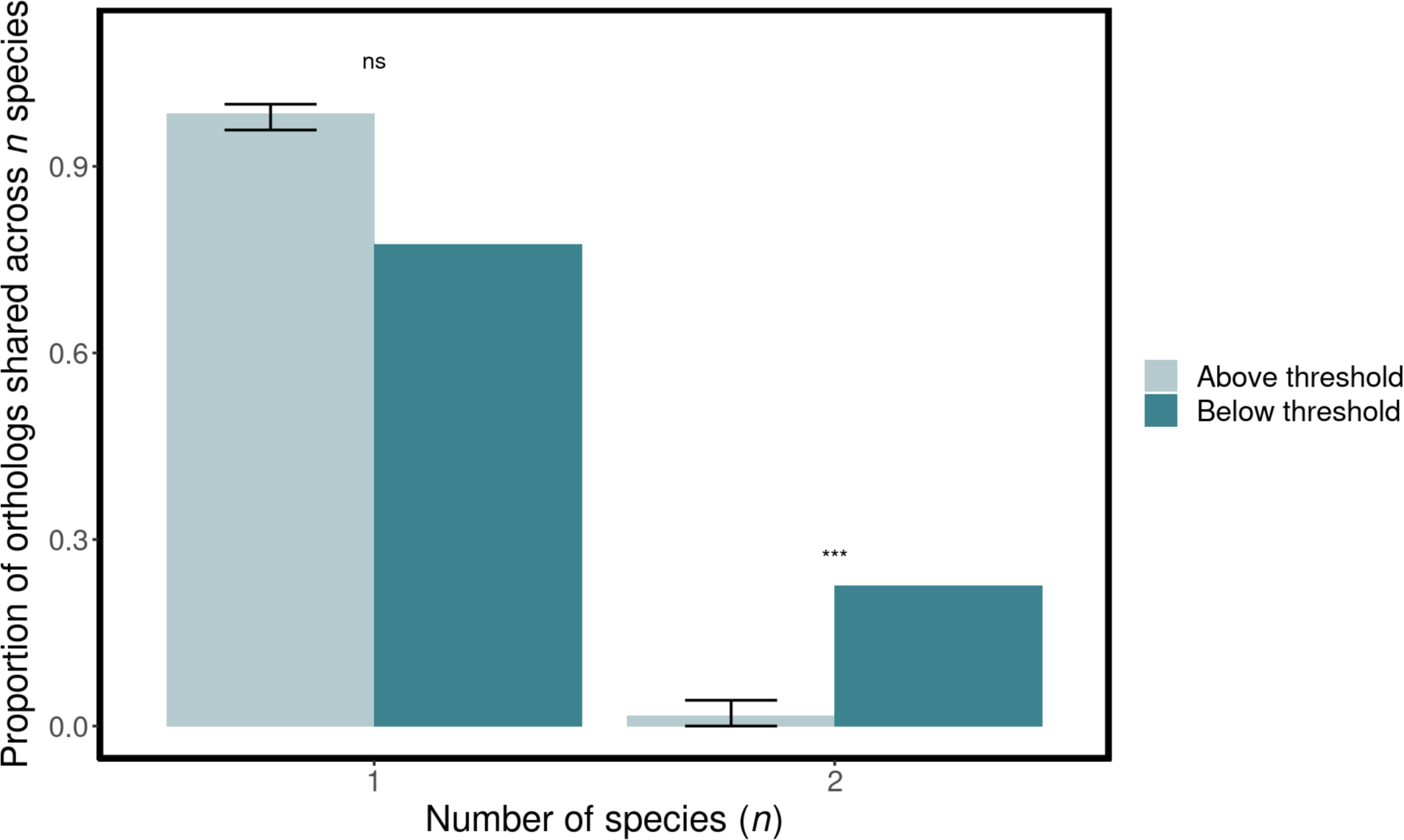
Proportion of orthologs shared across species in windows below and above the low diversity threshold in *D. melanogaster* (ZI) and *D. melanogaster* (RA) only. We obtained 100 random samples of orthologs found in windows above the low diversity threshold and obtained the mean number of orthologs shared across *D. melanogaster* populations (light blue) as well as the corresponding 95% confident intervals. We show the comparison between *D. melanogaster* populations in order to emphasize processes that may be influencing different populations of the same species.

**Fig S14.**
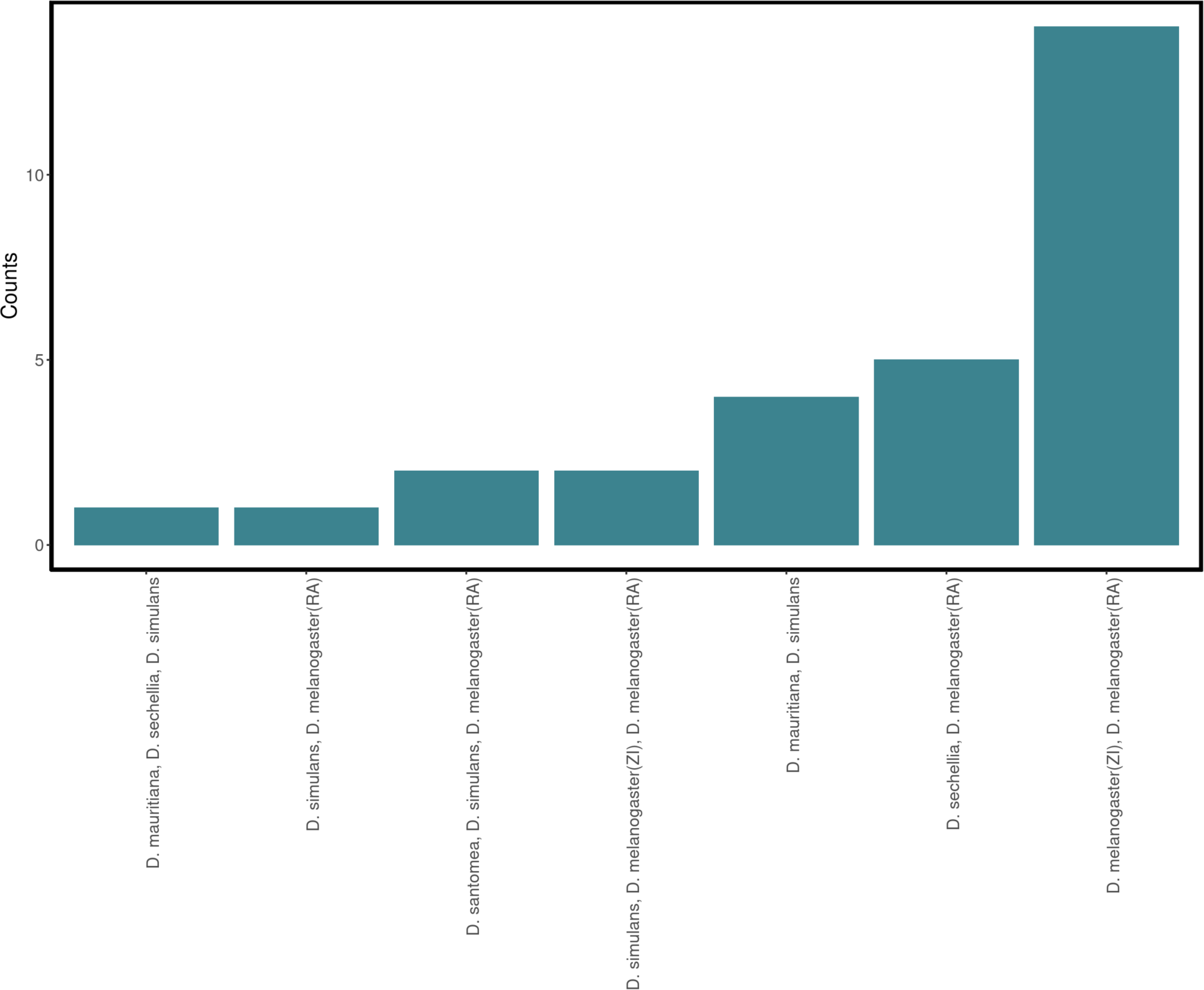
Number of orthologs shared across species. The x-axis shows the clusters of species that have orthologs in common. The *D. melanogaster* populations have the highest number of overlapping orthologs. Overlap between three or more species is less common.

## Text S1: Overlapping orthologous gene groups on the X chromosome across species

To investigate whether the low diversity regions observed across species on the X chromosome correspond to similar functions under selection across species, we identified orthologous gene groups present in two or more species for regions both below and above the low diversity threshold (**Methods**). We found that, excluding *D. melanogaster* (ZI) versus *D. melanogaster* (RA) comparisons, the proportion of orthologous genes that are present across three species was significantly higher in low diversity regions on the X compared to the rest of the chromosome (**Fig. S12**). We did not find a significant difference for orthologous genes found in two species. Specifically, we observed that 5% and 2% of orthologs in low diversity windows were shared across 2 or 3, respectively. In contrast, for regions above the threshold, 6% and 0.3% of orthologs where identical across 2 or 3 species, respectively. Moreover, when we included all populations, we found the highest overlap was between *D. melanogaster* (ZI) and *D. melanogaster* (RA) (14/ 43 orthologous gene groups found in at least two species; see **Fig. S13, Fig. S14**). This finding aligns with our expectations since *D. melanogaster* (ZI) and *D. melanogaster* (RA) are two populations of the same species, suggesting that the selection process may have originated in the ancestral population and persisted in the derived population. Despite the several instances of shared targets of selection across species, the majority of genes found in low-diversity regions were found in a single species (**Fig S12, Fig. S13**), indicating that distinct genes are targeted by selection independently.

